# Iron treatment induces defense responses and disease resistance against *Magnaporthe oryzae* in rice

**DOI:** 10.1101/2021.12.09.471912

**Authors:** Ferran Sánchez-Sanuy, Roberto Mateluna Cuadra, Kazunori Okada, Gian Attilio Sacchi, Sonia Campo, Blanca San Segundo

## Abstract

**Background:** Iron is an essential micronutrient required for plant growth and development. The impact of iron in plant-pathogen interactions is also well recognized. However, the molecular basis underlying the effect of plant iron status and immune function in plants is poorly understood. Here, we investigated the impact of treatment with high iron in rice immunity at the cellular and molecular level.

**Results:** We show that treatment with high iron confers resistance to infection by the blast fungus *M. oryzae* in rice. Histochemical staining of *M. oryzae*-infected leaves revealed that iron and Reactive Oxygen Species (ROS) accumulate at high levels in cells in the vicinity of the infection site. During pathogen infection, a stronger induction of defense-related genes occurs in leaves of iron-treated plants. Notably, a superinduction of phytoalexin biosynthetic genes, both diterpene phytoalexins and sakuranetin, is observed in iron-treated plants during pathogen infection. As a consequence, phytoalexin accumulation was higher in iron-treated plants compared with control plants. Transcriptional alterations of iron homeostasis-related genes and a reduction in apoplastic iron content were observed in leaves of Fe-treated rice plants.

**Conclusions:** These results illustrate that the iron status plays a key role in the response of rice plants to pathogen infection, while reinforcing the notion that iron signaling and defense signaling must operate in a coordinated manner in controlling disease resistance in plants. This information provides a basis to better understand the molecular mechanisms involved in rice immunity.

## INTRODUCTION

Iron (Fe) is an essential element required for a wide range of biological functions during plant growth and development. Plants require Fe for photosynthesis, mitochondrial respiration and hormone biosynthesis, among other processes. Fe is associated to cellular reduction-oxidation reactions (redox reactions) and electron transfer chains. It is also a co-factor for a variety of proteins mediating redox reactions. Fe deficiency results in chlorosis, poor growth and reduced yields (Hänsch *et al.* 2009). Although Fe is a constituent of approximately 5% of Earth’s crust, the availability of Fe to plants is usually very low (Connorton *et al.* 2017). Therefore Fe deficiency is a common problem in agricultural systems worldwide. Fe bioavailability is limited in aerobic conditions and high pH soil conditions where Fe, is predominantly found in the form of insoluble ferric form (Fe^3+^), forming hydroxide complexes poorly available for living organisms. In contrast, under reductive conditions, Fe is present as the soluble ferrous form (Fe^2+^) (Connorton *et al.* 2017).

Rice has evolved two distinct strategies to take up Fe from the soil, referred to as Strategy I and Strategy II (Bandyopadhyay and Prasad 2021). Dicotyledonous and non-graminaceous monocotyledonous species use **Strategy I**, or **reduction strategy** to increase Fe^3+^ solubility. It involves the secretion of phenolics and soil acidification by the release of protons into the rhizosphere, and is mediated by Phenolics Efflux Zero 1 (PEZ1) and H^+^-ATPase, respectively. Iron is subsequently reduced from the ferric (Fe^3+^) form to a more soluble ferrous form (Fe^2+^) by a ferric reductase-oxidase (FRO) and then it is transported across the plasma membrane into the root cells by Iron-Regulated Transporter (IRT1, IRT2). The **Strategy II**, also called the **chelation strategy**, is used by graminaceous species. Here, roots secrete Fe^3+^ chelators of the mugineic acid (MA) family called phytosiderophores (PS). This pathway implies the biosynthesis of the non-proteinaceaous amino-acid nicotianamine (NA) from S-adenosyl-L-methionine by NA synthase (NAS) and then NA is synthesized into 2’-deoxymugineic acid (DMA), the precursor of MAs, by NA aminotransferase (NAAT), and DMA synthase (DMAS). Secretion of MAs from rice roots to the rhizosphere is mediated by *OsTOM1* (Nozoye *et al.* 2011).

Rice is typically cultivated under anaerobic conditions (paddy fields) where abundant Fe^2+^ is readily available to the plant, especially in soils with a low pH in which Fe^3+^ is reduced to the more soluble ferrous ion Fe^2+^. Under such conditions, unlike other graminaceous plants, the absorption of Fe^2+^ by the rice roots might cause severe Fe toxicity. Thus, excessive accumulation of Fe^2+^ can be harmful to the plant (Connorton *et al.* 2017; Schmidt *et al.* 2020). The formation of deleterious reactive oxygen species (ROS) through redox reactions between ferric (Fe^3+^) and ferrous (Fe^2+^) (Fenton reaction) leads to the oxidation of biomolecules (lipids, proteins, DNA) and damage to cellular structures, and eventually, cell death (Turhadi *et al.* 2019). Therefore, Fe acquisition, use and storage must be tightly regulated at the cell and tissue level to provide enough amounts for the plant metabolism while preventing accumulations of deleterious ferrous Fe. Organelles are crucial compartments for Fe storage and sequestration within the plant cell. In particular, vacuoles play a major role in accumulating Fe excess, and releasing Fe into the cytosol, if required for metabolism (Kar and Panda 2020). Also, plant ferritins serve to store Fe and for protection against Fe-catalyzed ROS production (Arosio *et al.* 2009). The storage capacity of plant ferritins (e.g *Atfer1-4*, *Osfer1-2*) is of aprox. 4500 Fe atoms in the bioavailable and non-toxic form of Fe (Briat *et al.* 2010). Plants regulate Fe concentration in different subcellular compartments in a homeostatic way through dynamic processes to avoid Fe toxicity (Müller *et al.* 2015).

Because Fe participates in the generation of ROS, and ROS production is generally associated with resistance to pathogen infection, it is not surprising that Fe availability affects the outcome of plant pathogen interactions (Aznar *et al.* 2015; Dangol *et al.* 2019; Herlihy *et al.* 2020; Liu *et al.* 2007; Torres *et al.* 2005; Ye *et al.* 2014). As an example, resistance to the necrotrophic pathogens *Dickeya dadantii* and *Botrytis cinerea* was observed in Fe-starved *Arabidopsis* plants supporting that crosstalk between Fe deficiency response and immunity occurs in Arabidopsis (Kieu *et al.* 2012). Fe-starved maize plants were found to be unable to produce ROS in response to *Colletotrichum* infection which correlated with increased susceptibility to this fungal pathogen (Ye *et al.* 2014). However, our understanding on the regulatory mechanisms that mediate plant Fe homeostasis and plant immunity is still limited.

During pathogen infection, there is a competition between the host and the pathogen for Fe as the pathogen must acquire this vital element from host tissues, while the plant might interfere with Fe acquisition as a defense strategy during infection. On the one side, plant pathogens employ diverse strategies for Fe acquisition from the host plant (i.e. secretion of high-affinity Fe-binding siderophores) (Aznar *et al.* 2015). On the other side, plants have evolved mechanisms to sequester iron from pathogens during infection, a phenomenum that was originally described in animals, the so called “nutritional immunity” (Weinberg 1975). The host plant might also capitalize the toxicity of iron through local accumulation of Fe leading to activation of an oxidative burst that can be toxic for the invading pathogen (Aznar *et al.* 2015).

Rice ranks among the 10 most important crops worldwide, with greater importance for developing countries in terms of food security and alleviation of malnutrition. The rice blast disease caused by the ascomycete fungus *Magnaporthe oryzae* is the most devastating fungal disease of cultivated rice worldwide (Dean *et al.* 2012; Fernandez and Orth 2018; Wilson and Talbot 2009). *M. oryzae* is an ascomycete fungus with a hemibiotrophic lifestyle that involves initial proliferation inside living host cells before switching to a destructive necrotrophic mode (Fernandez and Orth 2018; Wilson and Talbot 2009). Many studies have been carried out during the last years to elucidate the molecular and cellular mechanisms implicated in the rice response to *M. oryzae* infection (Fernandez and Orth 2018). In previous studies, we reported that Fe supply positively impacts blast disease resistance in hydroponically-grown rice plants (Peris-peris *et al.* 2017) supporting crosstalk between iron signaling and immune signaling in rice.

In this work, we investigated the effect of treatment with high Fe on resistance to infection by *M. oryzae* in rice at the molecular and cellular level. Histological staining of *M. oryzae*-infected leaves revealed ROS and Fe accumulation in cells which are located in the vicinity of the infection sites. Treatment with high Fe increased resistance to *M. oryzae* which was accompanied by stronger induction of defense gene expression. Fe-treated rice plants also showed a higher induction of phytoalexin biosynthesis genes during pathogen infection, which correlated well with accumulation of major rice phytoalexins in the infected leaves, both diterpenoid phytoalexins and sakuranetin. *M. oryzae* infection also causes transcriptional regulation of genes involved in Fe homeostasis in rice leaves which was accompanied by alterations in total Fe and apoplastic Fe content. Collectively, these results further support links between the Fe status in the rice plant and resistance to infection by the blast fungus *M. oryzae*.

## RESULTS

### Treatment with high Fe enhances resistance to M. oryzae infection in rice plants

Resistance to infection by *M. oryzae* was investigated in soil-grown rice plants that have been treated with high Fe. For this, the rice plants were grown under sufficient Fe supply (50 μM Fe) for 16 days. Then, half of the plants continued growth under sufficient Fe while the other plants were supplied with 1 mM Fe (henceforth control and high-Fe plants, respectively). Two different periods of treatment with high-Fe were used, 5 and 19 days. After 5 days of treatment, no phenotypic differences were observed between control and high-Fe plants (**Supplemental Figure S1A**). Root and leaf biomass (fresh weight), as well as chlorophyll content did not differ between control and high-Fe plants (**Supplemental Figure S1B**). To note, at this time of treatment, the Fe level in tissues of high-Fe were higher than that in control plants (root, stem and leaves) (**Supplemental Figure S1C**). High Fe treatment caused a significant down-regulation in the expression of typical iron deficiency-inducible genes (*OsIRO2, OsNAS1, OsNAS2*) in the rice roots, thus, supporting that the plant perceives and respond to Fe treatment (**Supplemental Figure S1D)**. When Fe treatment was carried out for a longer period of time (e.g. 19 days of treatment with 1 mM Fe), the plants developed symptoms of Fe toxicity in leaves (**Supplemental Figure S2A**). At 19 days of treatment, the high-Fe plants also showed a significant reduction in root and leaf biomass (fresh weight) and chlorophyll content compared control plants (**Supplemental Figure S2B**). To avoid that toxic effects caused by Fe accumulation could confound our results on disease resistance, a period of 5 days of treatment with high Fe (1mM Fe) was chosen in this work. Under these experimental conditions, no penalty on plant growth was observed.

Blast resistance was assayed on high-Fe (after 5 days of treatment) and control Fe plants. Compared with control plants, high-Fe plants consistently exhibited higher resistance to *M. oryzae* infection as revealed by visual inspection of disease symptoms, quantification of diseased leaf area and measurement of fungal biomass in pathogen-infected leaves (**Figure 1A**). These results were substantially similar to those previous reported in hydroponically-grown rice plants (Peris-peris *et al.* 2017). Thus, treatment with high Fe enhances resistance to *M. oryzae* infection in rice plants.

**Figure 1.**
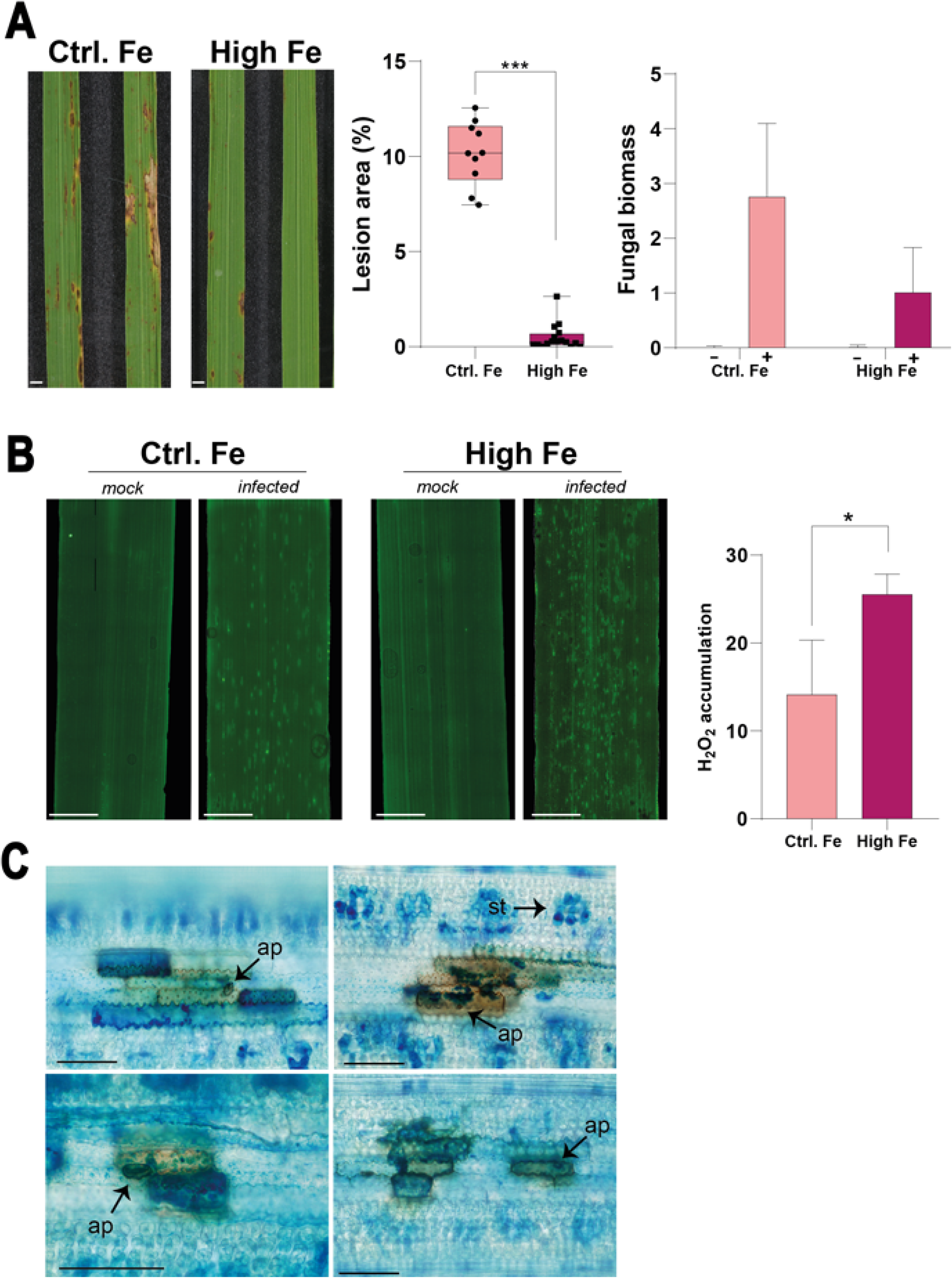
Resistance to *M. oryzae* infection in rice plants that have been grown under high Fe supply. Rice plants were grown in soil for 16 days under control Fe supply (0.05 mM Fe) and then supplied with high Fe (1 mM Fe; control plants) for 5 days more. Control plants were allowed to continue growth under control Fe supply. Rice plants were inoculated with a conidial suspension of *M. oryzae* spores (5 × 10^5^ spores/ml), or mock-inoculated. Data from one representative experiment of four independent experiments are presented. (**A)** Disease symptoms at 7 days post-inoculation (dpi). Right panels, percentage of diseased area at 7 dpi (n=10). Fungal biomass was quantification by qPCR using specific primers of the *M. oryzae 28S* ribosomal gene (relative to the rice *Ubiquitin 1* gene Os06g46770) at 7 dpi, (mock (-) and *M. oryzae* (+). Data are mean ± SEM (n=10). Asterisks indicate statistical significant differences calculated by *t*-test (*** indicate p < 0.001). (**B)** ROS accumulation in control and high-Fe rice plants under non-infection and infection conditions. ROS (H_2_O_2_) was detected using the fluorescent probe H_2_DCFDA at 48 hpi in mock-inoculated (mock) and *M. oryzae*-inoculated (infected) plants. Bars correspond to 2 mm. Right panel, Image J software was used for quantification of ROS fluorescence of three independent experiments (100 fields, each). Asterisks indicate statistical significant differences calculated by two-way ANOVA (*p < 0.05) (**C)** Accumulation of ROS (in brown) and iron (Ferric ions, Fe^3+^, in blue) in *M. oryzae*-inoculated leaves of high-Fe plants. The third leaf was stained with DAB (for ROS staining) followed by Prussian Blue (Perls reagent, for Fe staining) at 24-48 hpi. Bars correspond to 50 μm. Ap: apressorium; St, stomata.

A generalized plant defense against pathogen attack is the production of reactive oxygen species (ROS) (Torres 2010). Among ROS, H_2_O_2_ accumulation provokes localized cell death around the site of infection to limit the spread of the pathogen. Indeed, H_2_O_2_ is considered an important molecule in regulating plant immune responses (Torres *et al.* 2006). On this basis, we investigated whether treatment of rice plants with high Fe has an effect on ROS accumulation potentially contributing to the observed phenotype of blast resistance. Histochemical detection of ROS, mainly H_2_O_2,_ was carried out using the fluorescent probe H_2_DCFDA (2’, 7’ dichlorofluorescein diacetate) (Fichman *et al.* 2019) in leaves of control and high-Fe plants inoculated, or not, with *M. oryzae* spores. In the absence of pathogen infection, ROS accumulation was barely detected in control and high-Fe plants (**Figure 1B**, mock). Upon pathogen challenge, discrete regions accumulating ROS (H_2_O_2_) could be observed in control and high-Fe plants, but the DCFDA-fluorescent signals were more intense and abundant in high-Fe plants relative to control plants (**Figure 1B,** infected). The regions accumulating ROS, most probably, correspond to the *M. oryzae* infection sites. Quantification of H_2_DCFDA fluorescence using ImageJ software confirmed higher accumulation of ROS in *M. oryzae*-infected leaves of high-Fe plants than in *M. oryzae*-infected leaves of control plants (**Figure 1B,** right panel).

Knowing that rice plants grown under high Fe supply showed ROS accumulation during *M. oryzae* infection, and that Fe catalyzes the Fenton reaction to produce ROS (Krohling *et al.* 2016), it was of interest to investigate the sites of Fe accumulation in *M. oryzae*-infected leaves in Fe-treated rice plants. For this, we performed double staining using DAB (3, 3’-diaminobenzidine) for ROS detection followed by Perls staining for Fe detection. The Perls reagent (potassium ferrozianide) has been widely used for histochemical detection of Fe^3+^ ions in plant tissues, based on the capability of Fe^3+^ to react with potassium ferrozianide to produce Prussian blue (Roschzttardtz *et al.* 2010, 2013). For this study, we focused on high-Fe plants, these plants showing resistance to *M. oryzae* infection. Double staining of *M. oryzae*-infected leaves of high-Fe plants revealed ROS (brown) accumulation at the invaded cells, whereas Fe^3+^ (blue) accumulated in the invaded cells as well as in contiguous cells (**Figure 1C**). In previous studies, Dangol *et al.* (2019) described that Fe^3+^ and ROS (e.g. H_2_O_2_) accumulate in rice leaves during infection with an avirulent strain of *M. oryzae*. In non-infected rice leaves, iron accumulates at stomata (**Supplemental Figure S3**; similar results were previously reported by Sánchez-Sanuy *et al.* 2019). Resistance to *M. oryzae* infection in Fe-treated rice plants can then be explained by a localized accumulation of Fe and H_2_O_2_ at the sites of pathogen penetration.

### High Fe supply alters the transcriptome of rice plants

Most research to date on the rice response to Fe supply focused on transcriptional regulation of gene expression in roots (Ogo *et al.* 2014). Relatively less is known about transcriptional alterations induced by Fe treatment on rice leaves. Even less is known about the impact of Fe treatment on immune responses of rice plants to pathogen infection. Accordingly, in this work we initially performed a comparative transcriptome analysis of leaves from control and high-Fe plants. Differentially expressed genes (DEGs) were identified based on significance level (FDR ≤ 0.05) and log_2_ fold change (FC) with a threshold of FC ≥ + 0.5 and FC ≤ - 0.5 for up-regulated and down-regulated genes, respectively. Using these criteria, only 48 genes were found to be up-regulated in high-Fe plants compared to control plants, while 191 genes were down-regulated (the full list of DEGs is presented in **Supplemental Table S1a**). To note, genes whose expression is typically induced by Fe deficiency in rice roots where found to be repressed in leaves of high-Fe plants relative to control plants (**Figure 2A, Supplemental Table S1b**). They included: *OsIMA1* and *OsIMA2* (*iron deficiency-inducible peptide-IRON MAN*), *OsIRO2* and *OsIRO3* (*iron-related transcription factor 2 and 3*), *OsHRZ1* and *OsHRZ2* (Iron-binding Haemerythrin RING ubiquitin ligase). Other genes involved in iron homeostasis that are down-regulated in leaves of high-Fe rice plants are those encoding the plasma membrane iron transporters *OsOPT7* (*iron-deficiency-regulated oligopeptide transporter 7*), *OsNRAMP1* (*Natural Resistance-Associated Macrophage Protein 1*) and *OsVMT* (a vacuolar mugineic acid transporter) (**Figure 2A**). *OsFER2* (*FERRITIN 2*) expression was induced in leaves of high-Fe plants (**Figure 2A**). Ferritins are the primary Fe-storage proteins (in the form of Fe^3+^) and contribute to protection of plants against *Fe*-induced oxidative stress.

**Figure 2.**
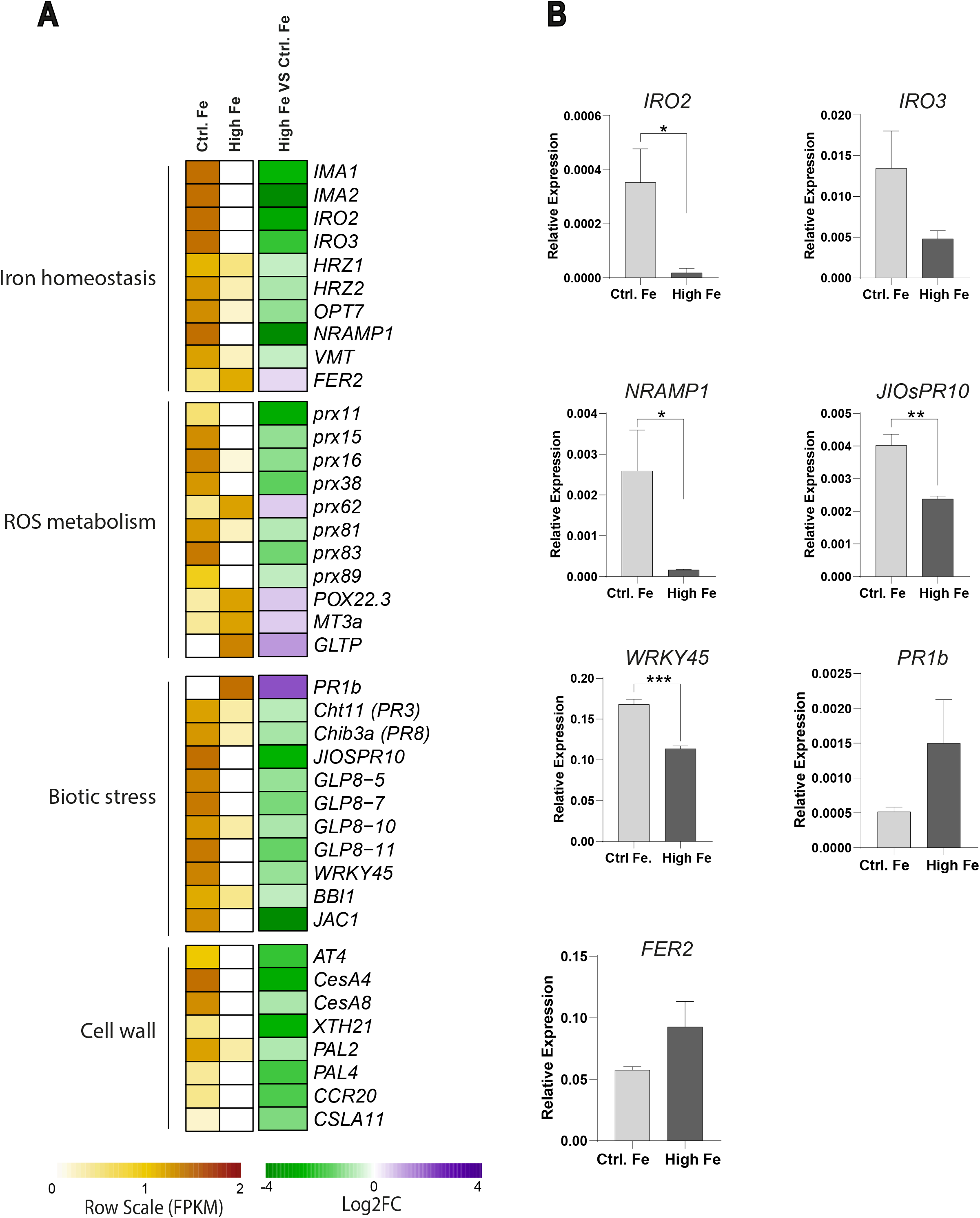
Differentially expressed genes in leaves of high-Fe plants relative to control plants. **(A).** Heatmaps showing differentially expressed genes (DEGs) by RNASeq analysis. Right panel, expression level (row scaled FPKM) is represented from pale yellow (less expressed) to brown (more expressed). Left panel, up-regulated genes (Log_2_ fold change (FC) ≥ + 0.5; purple) and down-regulated genes (Log_2_ FC ≥ - 0.5; green) DEGs. Data represented correspond to the mean of two biological replicates, each biological replicate consisting in a pool of 5 leaves from individual plants. The full gene name and ID of DEGs are indicated in **Supplemental Table S1a, S1b**. **(B)** Expression of differentially expressed genes identified by RNASeq analysis. Transcript levels were determined by RT-qPCR analysis in leaves of control and high-Fe plants. Data are mean ± SEM (n=3). Asterisks indicate statistical significant differences calculated by *t*-test (*, **, and *** indicate p < 0.05, 0.01, and 0.001, respectively). Gene-specific primers are listed in **Supplemental Table S8**.

Treatment with high Fe was also accompanied by alterations in the expression of genes implicated in detoxification and protection against oxidative stress. An important number of peroxidases were found to be misregulated by Fe treatment, which were either up-regulated (*OsPOX22.3, Prx62*) or down-regulated (*Prx11, Prx15, Prx16, Prx38, Prx81, Prx83, Prx89*) (**Figure 2A**). *OsPOX22.3* was found to accumulate in incompatible reactions of rice plants to *M. oryzae* (Faivre-Rampant *et al.* 2008). Besides catalyzing the decomposition of hydrogen peroxides, peroxidase enzymes have the capacity to produce H_2_O_2_, a process that is considered to be the source of H_2_O_2_ required for cell wall lignification. A Metallothionein-like protein (*OsMT3a*) and a glycolipid transfer protein involved in ceramide transport (*OsGLTP*) were induced also up-regulated in high-Fe plant compared with control plants (**Figure 2A**). Metallothioneins are metal-binding proteins implicated in metal detoxification and scavenging of ROS (Hassinen *et al.* 2011), while ceramides have been proposed to play a role in the induction of cell-death in rice, also linked to ROS accumulation (Zhang *et al.* 2020).

We also noticed that genes related to stress responses were misregulated genes in Fe-treated plants in the absence of pathogen infection, including genes for which a role in blast resistance has been described (**Figure 2A**). Among them, there were distinct members of several families of *Pathogenesis-Related* (*PR*) genes. Whereas *OsPR1b* was up-regulated in high-Fe plants compared with control plants, other *PR* genes were found to be down-regulated in these plants, such as *OsCht11* (PR3 family of PR proteins), *OsChib3a* (PR8 family), *JIOsPR10* (*Jasmonate Inducible PR10*; PR10 family) and four *OsGLP* genes (Germin-Like Proteins; PR14 family) (**Figure 2A**).

It is well known that the WRKY45 transcription factor plays a crucial role in resistance to *M. oryzae* infection, and that *WRKY45* overexpression confers resistance to *M. oryzae* (Shimono *et al.* 2012). Surprisingly, *OsWRKY45* was found to be down-regulated in high-Fe plants compared with control plants. Down-regulation of *PR* and *OsWRKY45* was in apparent contradiction with the phenotype of blast resistance that is observed in rice plants grown under high Fe conditions. Other defense-related genes down-regulated by Fe treatment were *OsBBI*, *OsJAC1* which are implicated in resistance to pathogen infection (Li *et al.* 2011; Weidenbach *et al.* 2016). Treatment with Fe also caused a general repression of key genes involved in plant cell wall biosynthesis (**Figure 2A**). Finally, RT-qPCR analysis confirmed RNA-Seq data (**Figure 2B**).

Together, these findings indicated that treatment with Fe was accompanied by alterations in the expression of genes that are typically activated by *M. oryzae* infection, even though the rice plants were not challenged with *M. oryzae*. Intriguingly, many of these genes showed down-regulated in high-Fe plants compared with control plants, although high-Fe plants exhibited enhanced resistance to *M. oryzae* infection. We then reasoned that blast resistance in high-Fe plants might rely more on the stronger induction of defense responses upon pathogen infection rather than constitutive defenses.

### High Fe supply boost defense-related gene expression during M. oryzae infection

We investigated pathogen-induced transcriptional changes in leaves of high-Fe and control plants. RNAs were obtained from *M. oryzae*-inoculated and mock-inoculated high-Fe and control plants at 48 hours post-inoculation (hpi). The same criteria described above (log_2_ FC ≥ 0.5 or ≤ −0.5, FDR ≤ 0.01, P ≤ 0.05) were used to examine the transcriptional response to *M. oryzae* infection in leaves of high-Fe and control plants compared with their respective non-infected controls. Pair-wise comparisons revealed 6470 pathogen-regulated genes in high-Fe plants (3293 up-regulated; 3177 down-regulated) (**Figure 3A, B; Supplemental Table S2a**). In control plants, 5996 genes were regulated by *M. oryzae* infection (2879 up-regulated; 3117 down-regulated) (**Figure 3A,B; Supplemental Table S2b**). Considering that the same criteria were used for identification of DEGs in both comparisons (e.g. infected *vs* mock in high-Fe and infected *vs* mock in control plants), this difference could be explained because pathogen infection causes stronger induction of those genes in high-Fe plants compared with control plants. In line with this, heat map visualization of RNASeq data showed stronger induction by pathogen infection in an important proportion of genes in high-Fe plants relative to control plants (**Supplemental Figure S4**).

**Figure 3.**
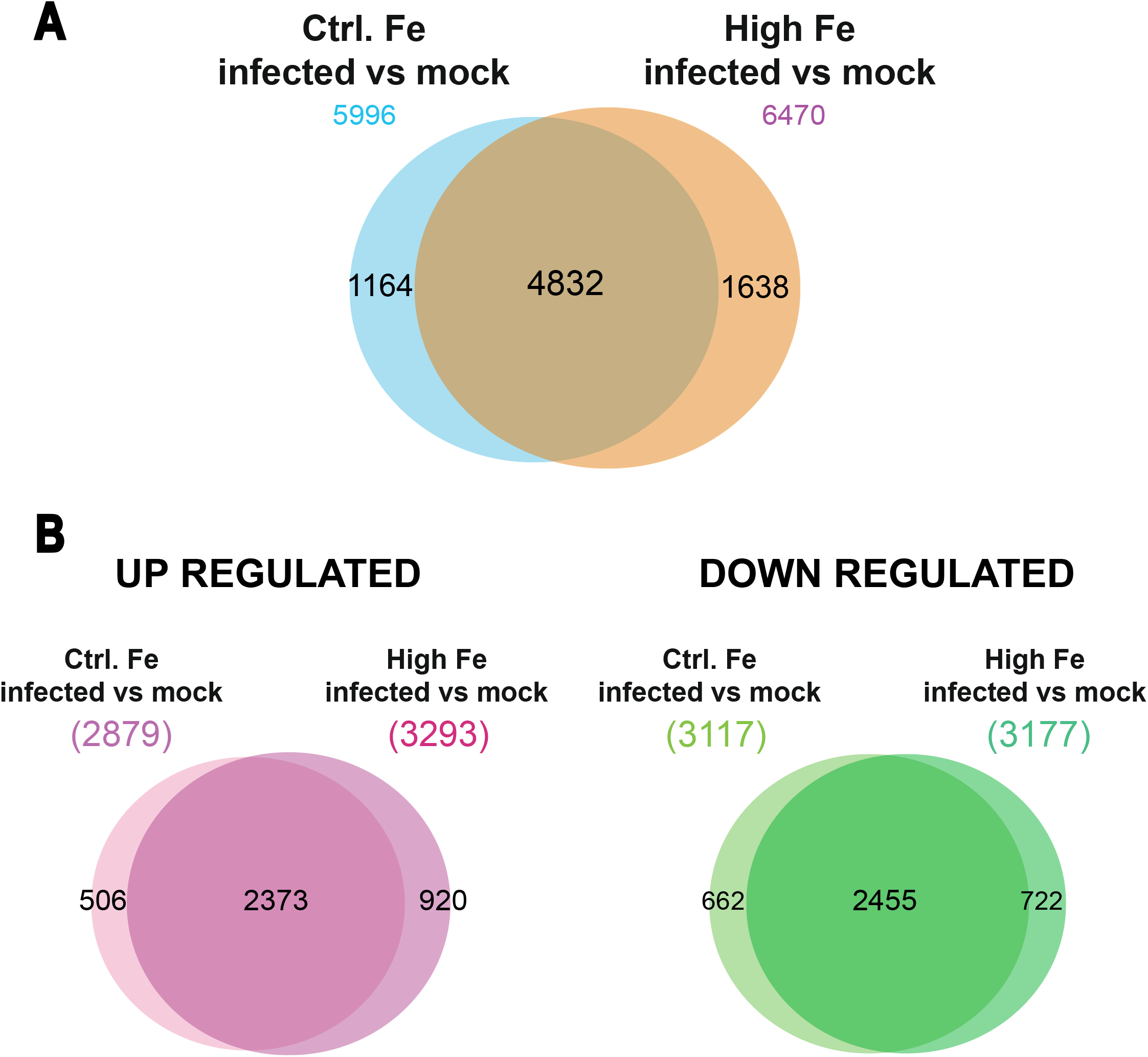
Response to *M. oryzae* infection in leaves of high-Fe and control plants. **(A)** Venn diagrams indicate the number of genes that are specifically and commonly regulated by *M. oryzae* infection in each Fe condition at 48 hpi. **(B)** Venn diagrams of up-regulated and down-regulated genes in each Fe condition (log_2_ fold change >0.5 or <-0.5; p-value ≤ 0.05).

Gene Ontology (GO) enrichment analysis was used to identify GO annotations in Biological Processes and Molecular Function associated to the response to *M. oryzae* infection in high-Fe and control plants. The REVIGO tool was used to remove redundant GO terms (Supek *et al.* 2011; http://revigo.irb.hr). Of interest, the most abundant subcategories in Biological Processes, in both high-Fe and control plants were “Diterpene Phytoalexin Metabolism” followed by “Secondary Metabolism” and “Defense Responses” (**Figure 4A; Supplemental Table S3A and S3B**). Another GO category highly represented in up-regulated was “Phosphorus Metabolism” (**Figure 4A**) which was consistent with the observed over-representation of the “ATP binding” and “Protein kinase activity” annotations in the Molecular Function category in high-Fe and control plants (**Figure 4B**). Also in the category of Molecular Function, genes involved in “Iron binding” and “Signaling receptor activity” were over-represented in genes that are up-regulated by infection in high-Fe and control plants, while genes with “Chitinase activity” were over-represented in high-Fe plants, but not in control plants (**Figure 4B**). Regarding genes down-regulated by *M. oryzae* infection in Biological processes, they categorized in “Photosynthesis” and “Pigment biosynthetic processes”, whereas in Molecular function, there were genes involved in “Iron-sulfur cluster binding” and “Zinc ion binding” (in both high-Fe and control conditions) (**Supplemental Figure S5**). Taken together, these results revealed similar GO terms associated to the response to *M. oryzae* infection in high-Fe and control rice plants.

**Figure 4.**
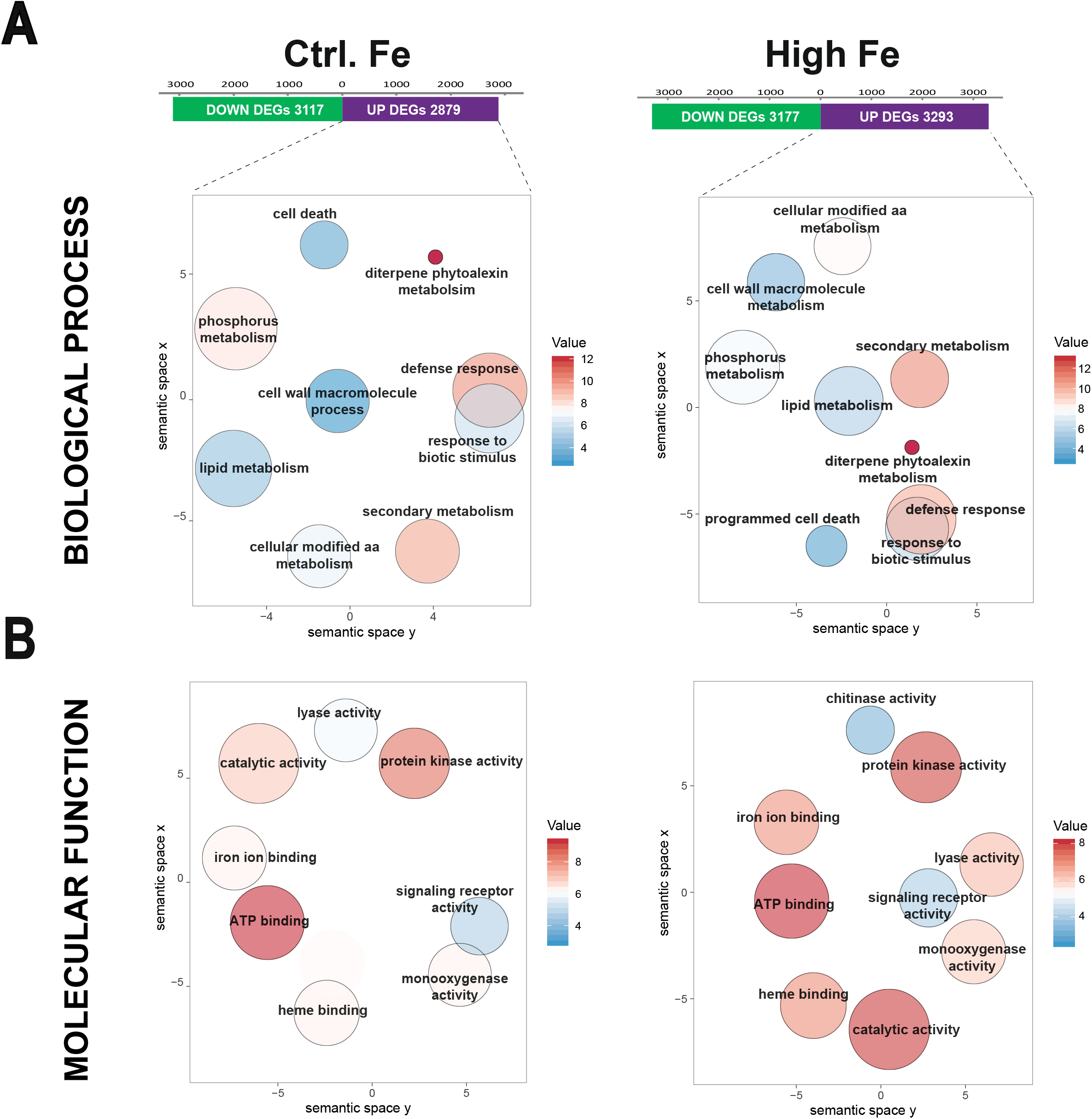
GO enrichment analyses of genes up-regulated by *M. oryzae* infection in control and high-Fe plants (48 hpi) in the categories of Biological Processes **(A)** and Molecular Function **(B)** GO terms were visualized using REVIGO (https://revigo.irb.hr/) after reducing redundancy and clustering of similar GO terms in the *O. sativa* database. GO terms are represented by circles and are clustered according to semantic similarities (more general terms are represented by larger size circles, and adjoining circles are most closely related). Circle size is proportional to the frequency of the GO term, whereas color indicates the enrichment derived from the AgriGO analysis (red higher, blue lower). Full data sets of DEGs and lists of GO terms are presented in **Supplemental Table S2 and S3**, respectively).

### Enhanced defense responses in iron-treated rice plants

Pathogen-induced changes in defense-related gene expression in high-Fe and control rice plants were compared. As expected, *M. oryzae* infection activated the expression of *PR* genes belonging to different *PR* families in both high-Fe and control plants (**Figure 5A; Supplemental Table S4**). They included *PR1, PR2* (β-1,3 glucanases)*, PR3, PR4* and *PR8* (chitinases)*, PR5* (thaumatin-like proteins, TLPs)*, PR6* (proteinase inhibitors)*, PR9* (peroxidases)*, PR10* (ribonuclease-like proteins), *PR14* (lipid transfer proteins, LTPs) and *PR15* (germin-like proteins). Among them, *OsPR1* and *OsPBZ1* (*PR10* family) are routinely used as marker genes for the induction of the rice response to *M. oryzae* infection (Agrawal *et al.* 2001; Midoh and Iwata 1996). Of interest, upon pathogen infection, *PR* genes were found to be activated to a greater extent in high-Fe plants compared with control plants (**Figure 5A; Supplemental Table S4**). It is well documented that PR proteins play an important role in plant defense against pathogen infection, and that overexpression of *PR* genes confers resistance to *M. oryzae* infection in rice (Jain and Khurana 2018; Wu *et al.* 2016). The antimicrobial activity of several PR proteins has been demonstrated (i.e. chitinases, thaumatins, LTPs, among others). Results obtained by RNA-Seq analysis were validated by qRT-PCR analysis of selected *PR* genes (*OsPR1a*, *OsPR1b*, *OsPBZ1,* a *PR10* family member, JI*OsPR10,* and *ROsPR10*) (**Supplemental Figure S6**). Thus, stronger induction of *PR* genes appears to occur in high-Fe plants compared with control plants.

**Figure 5.**
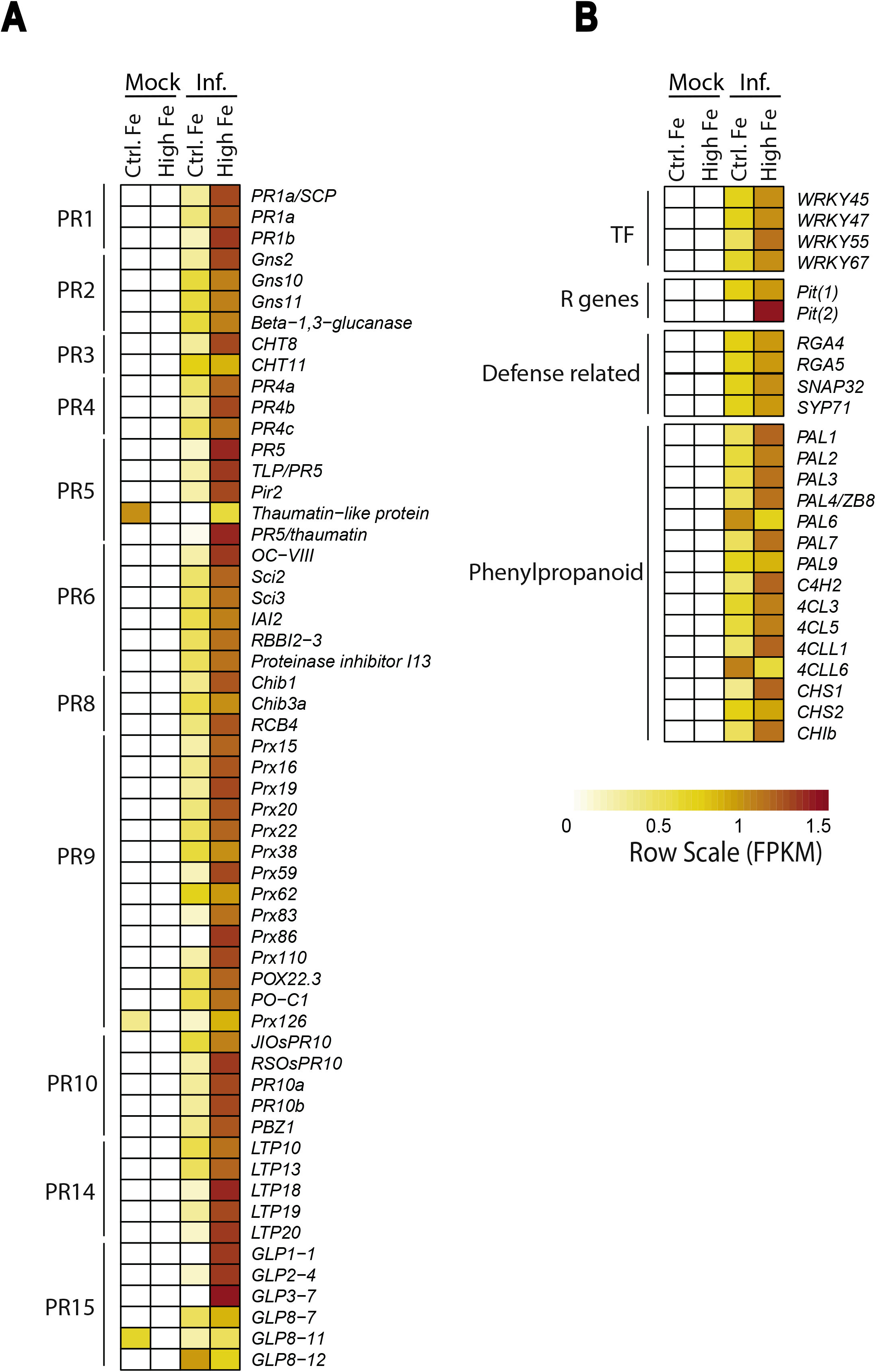
Expression of genes involved in the rice defense response to pathogen infection in leaves of control and high-Fe plants (48 hpi). Heat maps showing the expression level of DEGs. Left panel, gene expression is represented as row scaled FPKM from pale yellow (less expressed) to brown (more expressed). **(A)** Expression of *Pathogenesis-Related* (*PR*) genes. **(B)** Expression of defense-related genes. TF, transcription factors, R, Resistance genes. The gene name, FPKM and fold change values are indicated in **Supplemental Table S4**.

RNA-Seq analysis also revealed regulation of peroxidase genes by *M. oryzae* infection (up to 14 peroxidase genes; *Prx*, *PR9* family), most of them being induced at a higher level in high-Fe plants than in control plants (**Figure 5A; Supplemental Table S4**). Peroxidases catalyze oxidation of various substrates concomitant with the decomposition of H_2_O_2_. It is tempting to assume that *M. oryzae*-regulated peroxidases contribute to maintain proper H_2_O_2_ accumulation in high-Fe plants to avoid toxic effects to the plant cell. We also noticed that *OsWRKY45*, a positive regulator of blast resistance was induced at a higher level in high-Fe plants relative to control plants (**Figure 5B; Supplemental Table S4)**.

We also noticed that genes involved in the general phenylpropanoid pathway were induced to a different extent by Fe treatment; most of them being more strongly induced in high-Fe plants compared with control Fe plants. Among them, there were *phenylalanine ammonia lyase* (*PAL*), *cinnamic acid 4-hydroxylase* (*C4H*); *4-coumarate:CoA ligase* (*4CL*), as well as *chalcone synthase* (*CHS*) and *chalcone isomerase* (*CHI*) (**Figure 5B; Supplemental Table S4**). The steps in which these genes participate in the phenylpropanoid biosynthetic pathway are indicated in **Supplemental Figure S7**. Defensive functions of phenylpropanoid compounds have long been recognized, these metabolites conferring broad-spectrum disease resistance in different plant species (Yadav *et al.* 2020). Furthermore, treatment with high Fe was associated with stronger induction of diterpenoid phytoalexin genes (see below). Altogether, there results indicated that there is a correlation between the expression level of defense-related genes (e.g. higher induction in response to infection) and the increased resistance to *M. oryzae* that is observed in high-Fe plants.

### Effect of treatment with high Fe on the accumulation of rice phytoalexins

Phytoalexins are produced and accumulate in plants in response to pathogen infection, also in rice plants infected with the rice blast fungus *M. oryzae* (Duan *et al.* 2014). As previously mentioned, “Diterpenoid phytoalexin metabolism” was identified as the most enriched term in the set of genes that are up-regulated by *M. oryzae* infection in both high-Fe and control plants (see **Figure 4**). The major rice phytoalexins are diterpenoid phytoalexins and sakuretin, the only flavonoid phytoalexin so far identified in rice. Regarding diterpenoid phytoalexins, they are synthesized from the precursor molecule geranylgeranyl diphosphate (GGDP), the end product of the methylerythritol phosphate (MEP) pathway (**Figure 6A**). A heat map showing the expression of genes implicated in the MEP and diterpene phytoalexin biosynthesis pathways in response to *M. oryzae* infection in control and high-Fe plants is presented in **Figure 6B**. As expected, diterpenoid phytoalexin biosynthesis genes are not expressed in non-infected rice plants. Upon pathogen infection, the expression of genes involved in the production of major diterpene phytoalexins, namely oryzalexins A to F, phytocassanes A to E, momilactones A and B, and oryzalexin S was induced in both control and high-Fe plants. Notably, these genes were induced to a greater extend in high-Fe plants than in control plants (**Figure 6B; Supplemental Table S5**). Results obtained by RNA-Seq analysis were validated by qRT-PCR analysis of selected MEP and diterpenoid phytoalexin biosynthetic genes, namely *OsDXS3* (MEP pathway) and *OsCPS2, OsCPS4, OsMAS1, OsCYP701A8* and *OsCYP76M6* (diterpene phytoalexin pathway) (**Supplemental Figure S8**). Not only genes directly involved in diterpene phytoalexin biosynthesis, but also genes in the upstream methylerythritol phosphate (MEP) pathway leading to the precursor of diterpene phytoalexins GGDP (e.g. *OsDXS3*, *OsCMK*, *OsMCS* and *OsGGPPS1*) showed a higher induction in high-Fe plants compared with control plants (**Figure 6A, B; Supplemental Table S5**).

**Figure 6.**
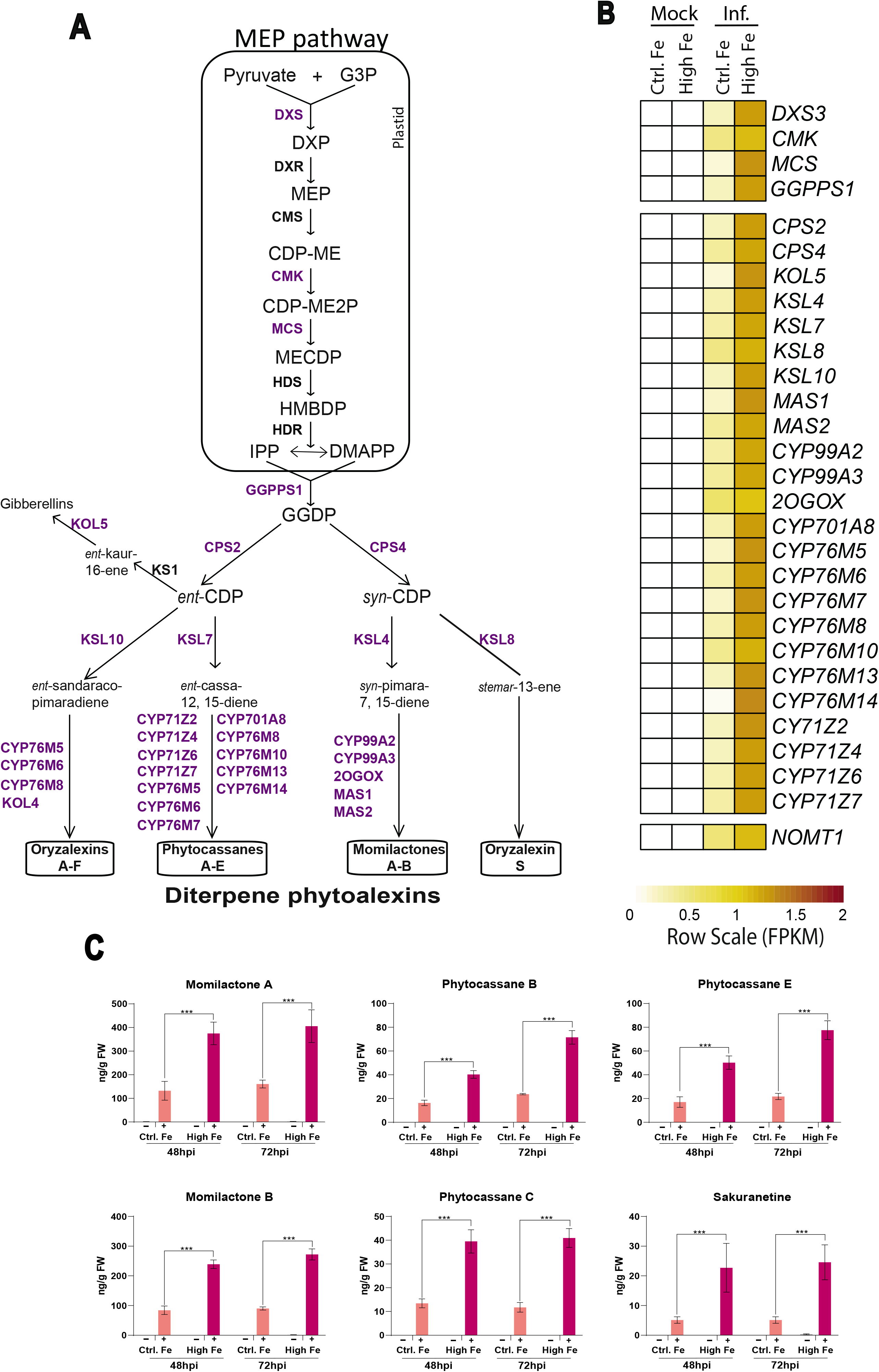
Expression of genes involved in phytoalexin biosynthesis and accumulation of phytoalexins in leaves of rice plants that have been grown under high Fe supply. **(A)** Methylerythritol phosphate (MEP) and diterpenoid phytoalexin biosynthesis pathways in rice. Genes whose expression is up-regulated by *M. oryzae* infection in control and high-Fe plants are indicated in purple color. The full name and details on the expression of these genes can be found in **Supplemental Table S5**. **(B)** Heat map showing the expression level (row scaled FPKM) at 48hpi. Left panel, gene expression is represented from pale yellow (less expressed) to brown (more expressed). **(C)** Accumulation of diterpenoid phytoalexins, momilactone A and B, phytocassane B, C, and E and the flavonoid phytohormone sakuretine in leaves of control and high-Fe plants, mock (-) and *M. oryzae*-infected (+) plants (48 and 72 hpi). FW, fresh weight. Four biological replicates (three technical replicates each) were analyzed. Data are mean ± SEM. Asterisks indicate statistical significant differences calculated by two-way ANOVA (*** p ≤ 0.001). **G3P** >, glyceraldehyde-3-phosphate; **DXP** >, 1-deoxy-D-xylulose 5-phosphate; **MEP** >, 2-C-methyl-D-erythritol 4-phosphate; **CDP-ME** >, 4-(cytidine 5’-diphospho)-2-C-methyl-D-erythritol; **CDP-ME2P**, 2-phospho-4-(cytidine 5’-diphospho)-2-C-methyl-D-erythritol; **MECDP**, 2-C-methyl-D-erythritol <2,4-cyclodiphosphate; **HMBDP**, 1-hydroxy-2-methyl-2-(E)-butenyl 4-diphosphate; **IPP**, isopentenyl diphosphate; **DMAPP**, dimethylallyl diphosphate; **GGDP**, geranylgeranyl diphosphate; and **CDP**, copalyl diphosphate.

As for the only flavonoid phytoalexin in rice, sakuranetin, it is synthesized from naringenin (phenylpropanoid pathway; see **Supplemental Figure S7**) by the activity of naringenin7-O-methyltransferase (NOMT). Our RNA-Seq analysis revealed higher expression of *OsNOMT1* in response to *M. oryzae* infection in high-Fe plants compared with control plants (**Figure 6**; **Supplemental Table S5**).

Knowing that the expression of phytoalexin biosynthesis genes was strongly induced by *M. oryzae* infection in high-Fe plants, it was of interest to examine phytoalexin content in these plants. As shown in **Figure 6C**, the accumulation of diterpenoid phytoalexins (Momilactone A and B, Phytocassane B, C and E) and sakuranetin drastically increased in *M. oryzae*-infected leaves. Phytoalexin accumulation was higher in high-Fe plants than in control plants, which is consistent with RNASeq data.

Together, these results indicated that blast resistance in high-Fe plants might result, as least in part, from supereractivation of phytoalexin biosynthesis genes, and subsequent accumulation of phytoalexins.

### Pathogen infection has an impact on the expression of iron homeostasis genes and iron content in rice leaves

The observation that *M. oryzae* infection provokes alterations in Fe^3+^ accumulation in distinct cells of the infected leaves (i.e. cells adjacent to the invaded cells; see **Figure 1C**) pointed to a regulation of genes involved in iron homeostasis in the *M. oryzae*-infected rice leaves. Indeed, RNASeq analysis revealed that a large number of genes with a known function in iron homeostasis are regulated by pathogen infection. Although many of these genes showed a similar trend in their response to infection in high-Fe and control plants, the intensity of the response varied depending on the Fe condition (e.g. stronger induction or repression by pathogen infection) (**Figure 7A,** control, infected vs mock; high-Fe, infected vs mock; **Supplemental Table S6**). Genes that are repressed by *M. oryzae* infection included genes encoding the transcription factors *OsIRO2*, *OsIRO3*, *OsIMA1*, *OsIMA2*, *OsHRZ1, OsHRZ2*, or the oligopeptide transporter *OsOPT7*, whose expression is induced in response to Fe deficiency (Bashir *et al.* 2015; Ogo *et al.* 2017; Kobayashi *et al.* 2013; Wang *et al.* 2020) (**Figure 7A; Supplemental Table S6**)

**Figure 7.**
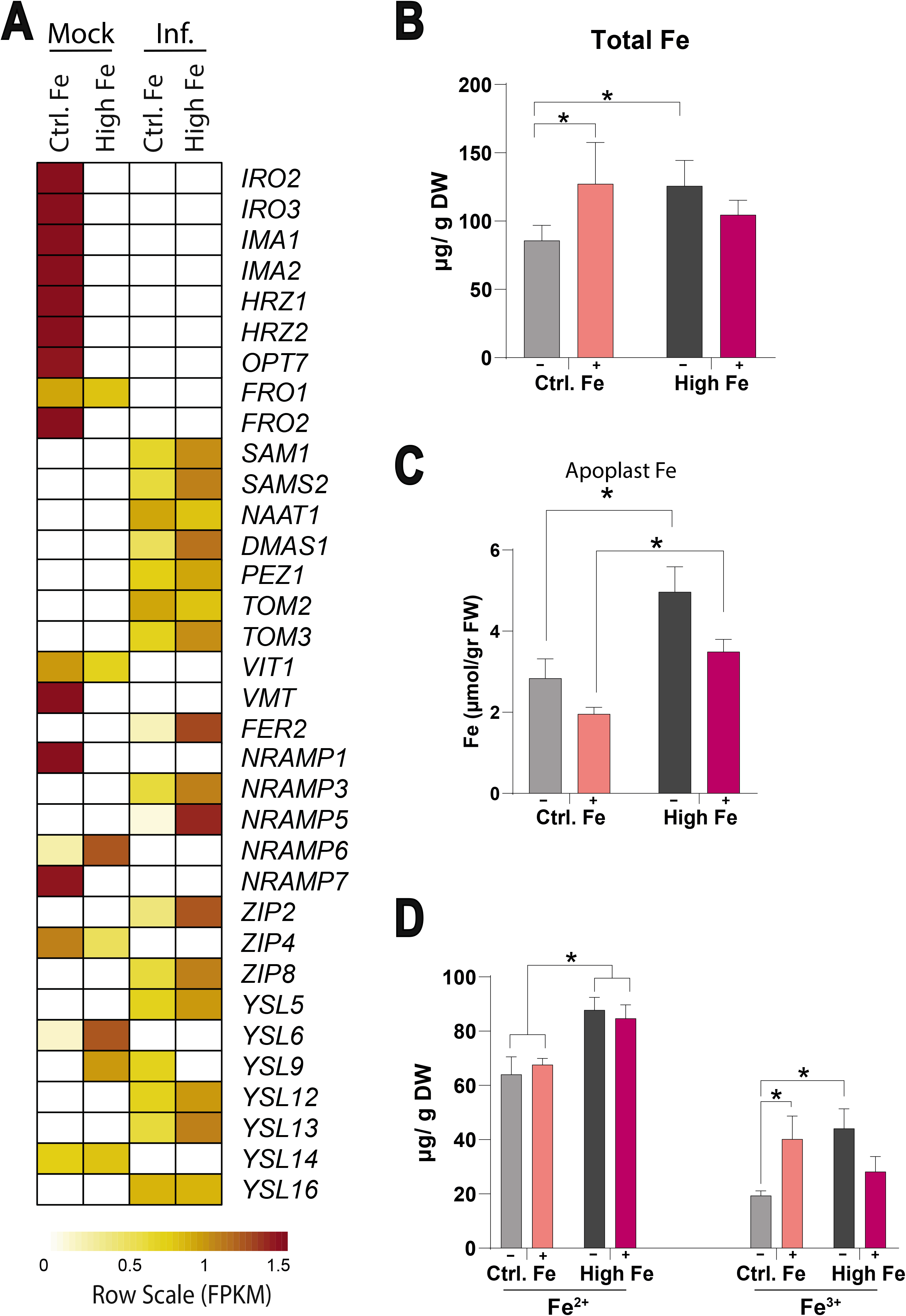
Expression of genes involved in Fe homeostasis and Fe accumulation in leaves of rice plants that have been grown under high-Fe supply. **(A)** Heat map showing the expression level (row scaled FPKMs) at 24 hpi. Left panel, gene expression is represented from pale yellow (less expressed) to brown (more expressed). The full name and details on the expression of these genes are indicated in **Supplemental Table S6.** Total Fe content estimated by colorimetric Ferrozine method in the youngest developed leaf (third leaf) at 48 hpi. **(C)** Fe content in the apoplast of leaves (youngest developed leaves) in *M. oryzae*-inoculated (+) and mock-inoculated (-) leaves. Fe content was estimated by ICP-MS. **(D)** Fe^2+^ and Fe^3+^ content was determined by the colorimetric Ferrozine method in the youngest developed leaf of control and high-Fe plants at 24 hpi. DW, dry weight. Five independent biological replicates (two technical replicates each) were analyzed in B, C and D. Data are mean ± SEM. Asterisks indicate statistical significant differences calculated by two-way ANOVA *, p ≤ 0.05).

As previously mentioned, excess Fe might become toxic to the plant via the Fenton reaction in which Fe^2+^ reacts with H_2_O_2_ to produce the hydroxyl radical (·OH), the most reactive ROS species. Although Fe^3+^ also reacts with H_2_O_2_ (Fenton-like reaction) to produce Fe^2+^ (as a source for subsequent production of toxic radicals in the Fenton reaction), the reaction of Fe^3+^ with H_2_O_2_ is substantially slower than the reaction of H_2_O_2_ with Fe^2+^. Inside the cell, the availability of free Fe^2+^ must be tightly controlled to avoid the Fenton’s reaction to occur. Storing Fe and/or Fe partitioning into sub-cellular compartments, mainly apoplast and vacuoles, are crucial to alleviate Fe toxicity (Moore *et al.* 2014). Ferric Reductase Oxidase (FRO) enzymes play an important role in maintaining proper Fe intracellular distribution and transport between the cytoplasm and the vacuole (Li *et al.* 2019). FRO is encoded by two genes in the rice genome (*OsFRO1* and *OsFRO2*). Of them, *OsFRO1* was reported to be responsible of the reduction of ferric Fe (Fe^3+^) into ferrous Fe (Fe^2+^) in the vacuole, which makes the vacuolar Fe available to the cytoplasm through Fe^2+^ transporters. Both *OsFRO1* and *OsFRO2* were found to be repressed by *M. oryzae* infection in control and high-Fe rice plants (**Figure 7**). Under high Fe conditions, downregulation of *OsFRO1* would reduce the amount of Fe^2+^ available for cytoplasm which would alleviate toxic effects caused by Fe excess. In the plant body, free Fe^2+^ is toxic; therefore, iron transport requires formation of complexes with phytosiderophores, such as the iron quelators of the mugineic acid (MA) family of phytosiderophores. To note, upon pathogen challenge, key genes related to the synthesis of MAs (*OsSAM1, OsSAMS2, OsDMAS1*) and phytosiderophore efflux transporters (*OsPEZ1, OsTOM3*) were induced at a higher level in high-Fe than in control plants.

On the other hand, *M. oryzae* infection strongly induced *FERRITIN2* (*FER2*) expression in high-Fe plants, and at a lesser extent also in control plants (**Figure 7A, Supplemental Figure S9**). Ferritin is the major iron-storage protein considered essential for tolerance to excess Fe. Its ability to sequester iron gives ferritin the function of protection against oxidative stress by mitigating damage from the Fenton reaction. In other studies, it was described that ferritins are up-regulated in various plant species following infection (Mata *et al.* 2001).

The NRAMP family of metal transporter proteins is of particular importance in iron compartmentalization which can be useful for Fe homeostasis, especially in conditions of excess of this element. RNASeq analysis showed that 4 genes encoding NRAMP proteins (Natural Resistance-Associated Macrophage Proteins) showed regulation by *M. oryzae* infection in control and high-Fe plants (**Figure 7A**). They were: *OsNramp1, OsNramp3, OsNramp5, OsNramp6 and OsNramp7* (all plasma membrane proteins). RT-qPCR analysis confirmed regulation of iron homeostasis-related genes in high-Fe and control plants *(OsIRO2, OsIRO3, OsFRO1, OsFRO2, OsNRAMP1, OsFER2)* (**Supplemental Figure S9**).

Other Fe-responsive genes whose expression is regulated by *M. oryzae* infection in rice leaves were members of the *Yellow Stripe-Like* (*YSL*) family of metal-NA (nonproteinogenic amino acid nicotianamine) transporters and zinc-regulated transporter ITR-like proteins (ZIP) family, including *OsYSL5*, *OsYSL6*, *OsYSL9*, *OsYSL12*, *OsYSL13*, *OsYSL14* and *OsYSL15* (among YSLs), and *OsZIP2*, *OsZIP4* and *OsZIP8* from the ZIP family (**Figure 7A**). NA was reported to be an indispensable component for the intracellular and intercellular distribution of iron in plants (Ishimaru *et al.* 2010). Regarding ZIP genes, although their expression is regulated by zinc (i.e. *OsZIP4)*, crosstalk between Fe and Zn homeostasis is known to occur in plants (Ishimaru *et al.* 2005).

Altogether RNASeq analysis revealed that *M. oryzae* infection triggers important transcriptional alterations in the expression of genes involved in Fe homeostasis. A differential regulation of these genes in high-Fe plants compared with control plants (e.g. stronger or weaker transcriptional responses) and spatial regulation of their expression in cells of the infected leaves might be crucial not only to arrest pathogen infection but also to alleviate Fe toxicity in the host plant.

Having established that pathogen infection regulates the expression of iron-associated genes, we examined Fe content in leaves of high-Fe and control plants, under infection and non-infection conditions (24 hpi). As expected, treatment with high Fe results in an increase in total Fe content (**Figure 7B)**. Upon pathogen challenge, there was a small but significant increase in total Fe content in leaves of control plants, but not in high-Fe plants (**Figure 7B**). To gain more information about effect of pathogen infection on iron content in rice leaves, we measured apoplastic Fe levels in *M.oryzae*-inoculated and mock-inoculated control and high-Fe plants. As shown in **Figure 7C**, *M. oryzae* infection is accompanied by a decrease in apoplastic Fe both in control and high-Fe plants, supporting perturbation of iron distribution in rice leaves during blast infection (e.g. cellular and apoplastic iron distribution). The concentration of apoplastic Fe in *M. oryzae*-infected high-Fe plants was similar to that in non-infected control plants.

Finally, since iron ions show unique reactivity depending on its redox state (e.g. Fe^2+^ or Fe^3+^), we investigated the oxidation states of Fe accumulating in *M. oryzae*-infected leaves of high-Fe and control plants which was then compared with that in the non-infected leaves for each condition. The ferrozine-based assay was used to quantify Fe^3+^ and Fe^2+^ content (Chen and Lewin 1972). In the absence of pathogen infection, treatment with high Fe increased both Fe^3+^ and Fe^2+^ levels (**Figure 7D**, grey and black bars). Upon pathogen challenge, Fe^2+^ content was not significantly altered in leaves of control or high-Fe plants (relative to the corresponding non-infected leaves). In the case of Fe^3+^, however, while pathogen infection increased Fe^3+^ content in control plants (infected *vs* non-infected control), its level decreased in high-Fe plants (infected *vs* non-infected high-Fe) (**Figure 7D**, pink bars). Thus, Fe^3+^ content in *M. oryzae*-infected high-Fe plants were similar to those in non-infected control plants.

Collectively, results obtained on the transcriptome analysis in combination with Fe analyses point to an important regulation in the expression of genes involved in iron homeostasis in rice leaves during *M. oryzae* infection. Treatment with high-Fe causes an increase in Fe content in rice leaves, total Fe (Fe^2+^ and Fe^3+^) and apoplastic Fe. Different response patterns are, however, observed between control and high-Fe plants during pathogen infection. Whereas in control plants pathogen infection increased the level of Fe (total Fe, and Fe^3+^), in *M. oryzae*-infected high-Fe plants Fe content decreased (total Fe, apoplastic Fe and Fe^3+^). A tight control of iron homeostatic mechanisms might occur during infection by *M. oryzae* in rice leaves.

## DISCUSSION

In this study, we provide evidence that the iron status of rice plants greatly influences resistance to *M. oryzae*. Several lines of evidence are consistent with this conclusion. First, we show that treatment with high Fe confers resistance to infection by the rice blast fungus. Second, blast resistance in high-Fe rice plants was associated to stronger induction of defense-related genes during pathogen infection, including *PR* genes. Third, a superinduction of genes involved in phytoalexin biosynthesis occurs during infection of high-Fe rice plants. Consequently, high-Fe rice plants accumulated major rice phytoalexins, diterpene phytoalexins (momilactones A, B, phytocassanes A-E, oryzalexins A-F and oryzalexin S), and sakuranetin. Diterpene phytoalexins and sakuretin are known to accumulate in rice leaves during *M. oryzae* infection, these metabolites exhibiting antifungal activity against *M. oryzae* (Hasegawa *et al.* 2014; Okada *et al.* 2007; Sánchez-Sanuy *et al.* 2019). The observed accumulation of phytoalexins might contribute, at least in part, to the phenotype of blast resistance in high-Fe plants.

Plants have evolved multifaceted strategies to avoid adverse consequences of Fe deficiency and excess in order to maintain normal growth and development. Consistent with an increase in total Fe content in Fe-treated plants, genes that typically induced by Fe deficiency were found to be repressed in high-Fe plants. Under the experimental conditions assayed in this work, however, Fe treatment had a low impact on the leaf transcriptome. Of interest, major differences between control and Fe-treated plants were observed during pathogen infection. Here, high-Fe plants developed a more robust defense response than control plants, which is in agreement with the phenotype of blast resistance that is observed in these plants. Superactivation of defense-related genes in high-Fe rice plants is reminiscent of defense priming, an adaptive strategy that improves the defensive capacity of plants (Gourbal *et al.* 2018; Martinez-Medina *et al.* 2016). In addition to *PR* genes and phytoalexin biosynthetic genes, our transcriptomic data showed stronger induction of *OsWRKY45* in high-Fe plants than control plants in response to *M. oryzae* infection. In other studies, an *OsWRKY45*-dependent priming of diterpenoid phytoalexin biosynthesis genes was described, while *OsWRKY45* overexpression conferred resistance to the rice blast fungus in rice (Akagi *et al.* 2014). Moreover, *OsFER2* was strongly up-regulated after pathogen infection in Fe-treated rice plants. Ferritin is known to bind excess Fe in a safe and bioavailable form to avoid cellular Fe toxicity (Aung and Masuda 2020). The observation that *Ferritin* is regulated by Fe treatment as well as by *M. oryzae* infection supports that plants have evolved mechanisms to sustain Fe homeostasis and immune reactions in concerted action. In other studies, ferritin was implicated in plant responses to pathogen infection (Dellagi *et al.* 2005; Herlihy *et al.* 2020).

Considering the outstanding importance of Fe in the generation of ROS, and that ROS mediate the induction of defense responses, we investigated Fe and ROS accumulation in high-Fe rice plants that have been inoculated with *M. oryzae* spores. Dual histochemical staining revealed that both Fe and H_2_O_2_ accumulate at the sites where pathogen penetration occurs. This localized accumulation of Fe might well trigger ROS production for the activation of immune responses to arrest pathogen invasion in high-Fe rice plants. The possibility of transfer of Fe from distal, non-infected leaf regions towards the infection sites should be also considered. If so, changes in the expression of iron homeostasis genes in the fungal-infected leaves would facilitate recruitment of Fe towards the infection sites. In addition to Fe recruitment, iron released from the first invaded dead cells might also provoke oxidative stress, thus contributing to immunity. The observed alterations in Fe distribution and subsequent accumulation of ROS in high-Fe plants is consistent with ferroptotic cell death at the sites of pathogen penetration, a response that was recently described in rice plants infected with an avirulent *M. oryzae* strain (Dangol *et al.* 2019; Distéfano *et al.* 2021). Dangol and collaborators (2019) also described that infection with a virulent strain of *M. oryzae* did not induce accumulation of Fe, thus, resulting in a phenotype of susceptibility. It is tempting to hypothesize that during infection of Fe-treated rice plants with a virulent *M. oryzae* isolate (present work), the rice plant develops a response similar to that in non-treated plants upon infection with an avirulent *M. oryzae* strain (Dangol *et al.* 2019), by accumulating Fe and ROS at the sites of infection.

Without infection, higher levels of Fe (total Fe, apoplastic Fe, Fe^2+^ and Fe^3+^) were observed in high-Fe plants compared with control plants. Upon pathogen challenge, total Fe increased in control plants while slightly decreasing in high-Fe plants. The observed reduction in total Fe content in high-Fe plants in response to pathogen infection might be the consequence of a reduction in apoplastic Fe and/or Fe^3+^ in these plants.

Being a foliar pathogen, *M. oryzae* entirely depends on the host tissue to acquire nutrients required for pathogen growth, including Fe. As both partners, host and pathogen, compete for Fe, a tight control over Fe homeostasis is of central importance in determining the outcome of the interaction. In this multifaceted process, the host plant might create conditions that restrict fungal proliferation in the plant tissue. Thus, the rice plant might deploy strategies to sequester Fe away from the pathogen during infection, a response that has been observed in other plant species (Herlihy *et al.* 2020). There is then the possibility that the observed reduction in total Fe and Fe^3+^ content that occurs in high-Fe plants during *M. oryzae* infection (which does not occur in fungal-infected control plants) might represent a limiting factor for fungal growth in high-Fe plants. Also, from results here presented, localized accumulation of Fe at the sites of pathogen infection, and subsequent accumulation of ROS, would provoke stronger induction of defense mechanisms in high-Fe rice plants, while increased ROS can be potentially toxic to the pathogen. On the other hand, the pathogen might use strategies for Fe acquisition from the host plant for its own nutritional needs, such as the production of siderophores (Hof *et al.* 2009). Also, when considering the influence that the Fe status in the host plant might have on the pathogen, it should be taken into account that the effect might depend on the pathogen lifestyle (necrotroph, biotroph, hemibiotroph pathogens). It will then be of great interest to investigate whether treatment with Fe confers resistance to pathogens with different lifestyles in rice.

Several studies have gathered evidence for a cross-talk between Fe signaling in Fe-starved plants and immunity (Aznar *et al.* 2015; Herlihy *et al.* 2020; Kieu *et al.* 2012; Verbon *et al.* 2017). For instance, Fe deficiency was found to confer resistance to *D. dadantii* and *B. cinerea* in Arabidopsis (Kieu *et al.* 2012). In other studies, Fe-starved maize was unable to produce ROS during infection with *Colletotrichum graminicola* which correlated with increased susceptibility to this fungal pathogen (Ye *et al.* 2014). These authors also described that maize recruits Fe^3+^ to the *Blumeria graminis* infection sites (Ye *et al.* 2014). Wheat plants infected with *B. graminis* f.sp. *tritici*, accumulated Fe^3+^ at cell wall appositions to mediate an oxidative burst (Liu *et al.* 2007). Results here presented on rice plants treated with high Fe, together with those previously reported by Dangol and collaborators (2019) on rice plants infected with an incompatible *M. oryzae* strain, demonstrated that Fe accumulates at the sites of fungal infection. From the information gained in maize, wheat and rice plants, it appears that accumulation of Fe, and subsequent ROS production, might be a critical response to arrest pathogen infection in cereal species. As most studies on the Fe nutritional status and immunity have been carried out in plants under Fe-limiting conditions, the main challenge now is to elucidate how iron homeostasis is controlled in leaves of rice plants under high Fe supply, and how infection by a foliar pathogen modulates these processes.

## CONCLUSIONS

In conclusion, the results presented here demonstrated that Fe plays a critical role in regulating immune responses in rice, and further supports cross-talk between Fe signaling and immunity in rice plants. Protection against *M. oryzae* infection can be explained by a localized accumulation of Fe at the sites of pathogen penetration. A tight regulation of Fe distribution in the infected leaves would then contribute to blast resistance. The mechanistic connection between accumulation of Fe and a successful immune response is explained at least in part by the iron’s capacity to produce ROS. A better understanding on the mechanisms underlying Fe homeostasis in leaves will allow the development of appropriate strategies for protection of rice against the blast fungus. Our observations provide an important basis for future investigation into the molecular mechanisms underlying Fe homeostasis in rice plants treated with high Fe.

## MATERIALS AND METHODS

### Plant and fungal material

Rice plants were grown at 28 °C with a 14 h/10 h light/ dark cycle. For Fe treatment, plants were grown in soil (turface:vermiculite:quartz sand [2:1:3]), and watered with a half-strength Hoagland solution during 14 days (2.5 mM KNO_3_, 2.5 mM Ca(NO_3_)_2_·4H_2_O, 1 mM MgSO_4_·7H_2_O, 0.5 mM NH_4_NO_3_, 25 μM KH_2_PO_4_, 23.15 μM H_3_BO_3_, 4.55 μM MnCl_2_·4H_2_O, 0.38 μM ZnSO_4_·7H_2_O, 0.1 μM CuSO_4_·5H_2_O, 0.14 μM Na_2_MoO_4_·2H_2_O, 50 μM Fe-EDDHA, pH 5.5). After 14 days, the plants were supplied with the same nutrient solution containing 1 mM Fe-EDDHA or maintained at 50 μM Fe-EDDHA. After 5 days of treatment, rice plants were infected with *M. oryzae* spores as described below.

### Blast resistance assays

The fungus *M. oryzae* (strain Guy-11) was grown in Complete Media Agar (CMA, 9 cm plates, containing 30 mg/L chloramphenicol) for 15 days at 28 °C under a 16 h/8 h light/dark photoperiod condition. *M. oryzae* spores were prepared as previously described (Campo *et al.* 2013). Soil-grown plants (3–4 leaf stage) were spray-inoculated with *M. oryzae* spores (5×10^5^ spores/ml; 0.3 ml/plant) using an aerograph at 2 atm of pressure. Plants were maintained overnight in the dark under high humidity, and allowed to continue growth for the required period of time. One of three independent experiments with similar results is shown (n=10).

### Phenotypical analysis and chlorophyll content

Chlorophyll content was determined using the SPAD 502 Plus Chlorophyll Meter (Spectrum Technologies). SPAD readings were obtained in the youngest developed leaf of rice plants grown in different Fe concentrations. Data are mean of 20 biological replicates. Shoot and root fresh weight (FW) was determined in three-week-old rice plants treated, or not, with 1 mM Fe at 5 days and 19 days after the onset of treatment. Data are mean of 10 biological replicates.

### Plant tissue staining

H_2_O_2_ determination was carried out using H_2_DCFDA (2’,7’-dichlorodihydrofluorescein diacetate) staining following the procedure described by Kaur *et al.* 2016. Briefly, the rice leaves were cut into pieces (3 cm) and incubated in a solution containing 10 μM H_2_DCFDA, vacuum infiltrated during 5 min, and then maintained in darkness at room temperature for 10 min. The samples were washed three times with distilled water and stored in 20% glycerol and immediately visualized. The fluorescence emitted by H_2_DCFDA was observed at 488 nm in an AixoPhot DP70 microscope under GFP fluorescence (488 nm excitation, 509 nm emission). Fluorescence was quantified using ImageJ software.

DAB (3,3 Diaminobenzidine) and Perls staining were carried out in mock- and *M. oryzae*-inoculated rice leaves, at 48 h post-inoculation [hpi]. For DAB staining, the rice leafs were cut in 3 cm pieces, incubated in a solution containing 1mg/ml DAB, vacuum infiltrated during 30 minutes, and then maintained in the dark at room temperature overnight. Leaves were washed with 75% ethanol.

Following DAB staining, the rice leaves were subjected to Perls staining following the procedure described by Stacey *et al.* (2008) with some modifications. Briefly, rice leaves were vacuum-infiltrated in a fixing solution (chloroform:methanol:glacial acetic acid; 6:3:1, v/v) for 1 h and incubated overnight at room temperature. After washing with distilled water (three times), samples were vacuum infiltrated with a pre-warmed (37 °C) staining solution (4% HCl and 4% K-ferrocyanide at equal volumes) for 1 h, incubated 4 h more at 37 °C in the same solution without vacuum, and then washed three times with distilled water. Sections were mounted in glycerol 50%. All the images were taken in an AixoPhot DP70 microscope under bright light. Three biological replicates with 4 technical replicates each were performed.

### RNA-seq analysis

Total RNAs were obtained using Maxwell(R) RSC Plant RNA Kit (Promega). For RNA-Seq, two biological replicates for each condition were examined, each biological replicate consisting of leaves from five individual plants (youngest totally expanded leaf). RNA concentration and purity were checked using a spectrophotometer (NanoDrop, ND-1000). RNA quality and integrity was examined on an Agilent 2100 Bioanalyzer (Agilent Technologies, Inc.; RNA integrity number (RIN) ≥ 8). A total of eight libraries were subjected to RNA-Seq analysis (125 paired-end reads), yielding an average of 24 812 610 clean reads/library. Raw reads were processed and analyzed as previously described (Sánchez-Sanuy *et al.* 2019). Reads were mapped against the reference genome, *Oryza sativa* sp. *japonica* (IRGSP-1.0 Ensembl release 41). To identify differentially expressed genes, a FDR cutoff < 0.01 and log2FC −0.5 ≤ or ≥ 0.5 was applied. Gene Ontology (GO) enrichment analysis (GOEA) was performed using the AgriGO web tool (Parametric Analysis of Gene Set Enrichment) (Du *et al.* 2010, https://bioinfo.cau.edu.cn/agriGOv2/). Enriched GO terms were clustered and plotted with the online analysis tool ReviGO (https://revigo.irb.hr/, Supek *et al.* 2011).

### Expression analysis by qRT-PCR

Total RNA was extracted from plant tissues by using TRizol reagent (Invitrogen). For quantitative RT-PCR (RT-qPCR), the first complementary DNA was synthesized from DNase-treated total RNA (0.5 μg) with High Capacity cDNA Reverse Transcription (Life technology, Applied Biosystems). Amplification involved cDNA (2 μl, 5 ng/μl) in optical 384-well plates (Roche Light Cycler 480; Roche Diagnostics, Mannheim, Germany) with SYBR Green I dye and gene-specific primers (**Supplemental Table S8**). The *Ubiquitin1* gene (Os06g0681400) was used to normalize transcript levels. Results from one of three independent experiments with similar results are shown. Three biological replicates and two technical replicates were analyzed. Asterisks indicate significant differences calculated by two way ANOVA *, p ≤ 0.05; **, p ≤ 0.01; ***, p ≤ 0.001).

### Analysis of apoplastic Fe

The apoplast wash fluid (AWF) was obtained from the same plant material used for the analysis of total Fe content as previously described (Kim *et al.*, 2013). Briefly, the youngest developed leaves of three-week-old plants (1 gr of fresh weight) were cut in 5 cm pieces and placed in a tube containing 30 ml of 200 mM CaCl_2_, 5 mM Na-acetate (pH 4.3 adjusted with glacial acetic acid) and 0.1 mM TPCK and 0.1 mM PMSF (freshly prepared), and kept on ice under constant agitation for 1 hour. Then, vacuum was applied for 30 minutes and leaves were removed and carefully dried. Dried leaves were rolled and placed in a tube for AWF recovery. Leaves were centrifuged a 3000 rpm, 15 minutes at 4°C. AWF was recovered from the bottom of the tube and kept in - 20 until use.

To assess that the apoplast fluid was devoid of cytoplasmic contaminants we performed the malate dehydrogenase (MDH) activity assay. For each material, MDH activities in total protein extracts was compared with that of apoplast samples. For this, total protein extracts were prepared from rice leaves using 1 mM MOPS, pH 7.5, containing 5 mM NaCl, and protease inhibitors (TPCK and PMSF, 0.1 mM each). Protein concentrations of total and apoplastic fluid extracts was quantified using Bradford. For MDH assay, protein samples (10 μl of either total protein or apoplastic fluid extracts, at 0.1 μg/μl) were added to the reaction mixture containing 0.4 mM NADH, 0.2 mM oxaloacetic acid, 1mM MOPS, pH 7.5, in 96 micro titer plates. The absorbance at 340nm at 25°C was recorded during 10 minutes. MDH activity was calculated as (Δabsorbance 340.min-1)/mg). Malate Dehydrogenase (MDH) activity was found to be in the range of 3.5% - 4% compared to total leaf protein extracts (**Supplemental Figure S10**).

Fe content was determined by inductively coupled plasma-mass spectrometry (ICP–MS; Bruker AURORA M90 ICP–MS, Bruker Daltonik GmbH, Bremen, Germany). Briefly, 30 μL of the apoplastic solutions were diluted to 1.2 mL with 2% HNO3 in (v/v) bidistilled water. In order to check the nebulization performance, an aliquot of 2 mg L-1 of an internal standard solution (72Ge, 89Y, and 159Tb) was added to the samples and calibration standards to give a final concentration of 20 μg L-1. Possible polyatomic interferences were removed by using CRI (Collision–Reaction-Interface) with an H_2_ flow of 80 mL min-1 flown though skimmer cone. Calibration curve were obtained using multi- and a single-(for P) ICP-MS standard solutions (Ultra Scientific, USA). Five biological replicates were analyzed. Asterisks indicate significant differences calculated by one way ANOVA *, P ≤ 0.05; **, P ≤ 0.01).

### Quantification of Ferrous (Fe^2+^) and Ferric Fe (Fe^3+^)

The ferrozine reagent was used for quantification of Fe^2+^. For Fe^3+^ quantification samples were treated with hydroxylamine, a reducing agent converting Fe^3+^ to Fe^2+^ (Stookey 1970). Ferrozine, in the presence of ferrous ions gives a pink-purple color which can be measured spectrophotometrically. Fe content was determined as previously described (Adolfsson *et al.* 2017). For this, the youngest totally expanded leaves of three-week-old plants were dried (5 days at 80° C). 50-70 mg of dried samples were pulverized in a Tissue Lyser, dried material was resuspended in 1M HCl (700 μl) and kept under constant agitation overnight. After centrifugation (10.000 rpm, 5 min, room temperature), the supernatant was recovered. Centrifugation was repeated twice. 150 μl were recovered for Fe^2+^ quantification and other 150 μl for total Fe quantification (Fe^2+^ + Fe^3+^). For Fe^2+^ quantification 15 μl of ferrozine 2 mM diluted in ammonium acetate 5M pH 9.5 were added to 150 μl of plant extract. For total Fe quantification 75 μl of hydroxylamine 6.25 M diluted in distilled water were added to the 150 μl of plant extract and incubated for 15 minutes. Then, 15 μl of ferrozine 2 mM diluted in ammonium acetate pH 9.5 were added. Blanks were carried out following the same procedures but adding 15 μl of ammonium acetate 5M pH 9.5 to plant extracts without ferrozine. Absorbance was read at 562 nanometers. Fe concentration was estimated by subtracting ferrozine based values with blank values. Fe^3+^ concentration was estimated by subtracting reduced hydroxylamine values (total Fe) with Fe^2+^ values. To determine final Fe concentration, standard curves at increasing Fe concentrations were performed. Five biological replicates with two technical replicates were performed. Asterisks indicate significant differences calculated by two way ANOVA *, p ≤ 0.05; **, p ≤ 0.01)

### Quantification of rice phytoalexins

For quantification of rice phytoalexins, leaf segments (aprox. 2 cm in length, 200–300 mg) were mixed with 40 vol. of 70% methanol and incubated at 4 °C overnight with constant rotation. A 1 ml aliquot was centrifuged at maximum speed to remove cell debris and used for quantification of phytoalexins.

Phytoalexins were quantified by liquid chromatography-tandem mass spectrometry (LC-MS/MS) as previously described (Miyamoto *et al.* 2016). Three biological replicates with two technical replicates each were performed. Significant differences in phytoalexin accumulation were evaluated with two way ANOVA *, p ≤ 0.05; **, p ≤ 0.01).

---

**Supplemental Figure S1.** Phenotype of rice plants that have been grown under control Fe (0.05 mM Fe, control plants) or high Fe (1 mM Fe, High-Fe plants) supply. Phenotypical parameters were assayed at 5 days of Fe treatment. **(A)** Appearance of control and high-Fe plants. (**B)** Shoot and root fresh weight (FW) (left and middle panels). Chlorophyll content (right panel). Differences were not statistically significant. **C** Total Fe content estimated by the ferrozine colorimetric method in roots, stem and leaf treated as in (A). Data are mean ± SEM (n=10). Asterisks indicate statistical significant differences calculated by *t*-test (*, **, and *** indicate p < 0.05, 0.01, and 0.001, respectively). **(D)** Expression of Fe deficiency responsive genes in roots of high-Fe and control plants after 5 days of Fe treatment. Transcript levels were determined by RT-qPCR analysis. The expression values were normalized to the rice *Ubiquitin1* gene. Three independent biological replicates (2 technical replicates each) were assayed. Data are mean ± SEM. Asterisks indicate statistical significant differences calculated by *t*-test (*, **, and *** indicate p < 0.05, 0.01, and 0.001, respectively). Gene-specific primers are listed in **Supplemental Table S8**.

**Supplemental Figure S2.** Phenotype of rice plants that have been grown under control Fe (0.05 mM Fe, control plants) or high Fe (1 mM Fe) supply. Phenotypical parameters were assayed at 19 days of treatment with high Fe. **(A)** Appearance of control and high-Fe plants. **(B)** Shoot and root fresh weight (FW) (left and middle panels). Chlorophyll content (right panel). Data are mean ± SEM (n=10). Asterisks indicate statistical significant differences calculated by *t*-test (* and *** indicate p < 0.05 and 0.001, respectively).

**Supplemental Figure S3.** Accumulation of iron (Ferric ions, Fe^3+^) in leaves of control and high-Fe rice plants. The third leaf was stained Prussian Blue (Perls reagent (blue color)). Bar, 50 μm.

**Supplemental Figure S4.** Heat map depicting the expression of genes (row scale FPKM) whose expression is regulated by *M. oryzae* infection in control and high-Fe rice plants. Right panel, expression level (row scaled FPKM) is represented from pale yellow (less expressed) to brown (more expressed). Left panel, up-regulated genes (Log_2_ fold change (FC) ≥ + 0.5; purple) and down-regulated genes (Log_2_ FC ≥ - 0.5; green) DEGs. Data represented correspond to the mean of two biological replicates, each biological replicate consisting in a pool of 5 leaves from individual plants.

**Supplemental Figure S5.** GO enrichment analyses of genes down-regulated by *M. oryzae* infection in control and high-Fe plants (48 hpi) in the categories of Biological Processes **(A)** and Molecular Function **(B)**. GO terms were visualized using REVIGO (https://revigo.irb.hr/) after reducing redundancy and clustering of similar GO terms in the *O. sativa* database. GO terms are represented by circles and are clustered according to semantic similarities (more general terms are represented by larger size circles, and adjoining circles are most closely related). Circle size is proportional to the frequency of the GO term, whereas color indicates the enrichment derived from the AgriGO analysis (red higher, blue lower). Full data sets of DEGs and lists of complete GO terms are presented in **Supplemental Table S2** and **S3,** respectively).

**Supplemental Figure S6.** Expression of *PR* genes in leaves of control and high-Fe plants (-, mock-inoculated; +, *M. oryzae*-inoculated). Transcript levels were determined by RT-qPCR analysis at 48 hpi. Expression values were normalized to the rice *Ubiquitin1* gene. Three independent experiments (2 technical replicates each) were carried out with similar results. Data are mean ± SEM (n=3). Asterisks indicate statistical significant differences calculated by two-way ANOVA (*, **, and *** indicate p < 0.05, 0.01, and 0.001, respectively). Gene-specific primers are listed in **Supplemental Table S8**.

**Supplemental Figure S7.** Phenylpropanoid biosynthesis pathway. **PAL**, phenylalanine ammonia lyase; **C4H**, cinnamate-4-hydroxylase; 4CL, 4-coumaroyl-CoA ligase; **CCR**, cinnamoyl-CoA reductase; **CHS**, chalcone synthase; **CHI**, chalcone isomerase; **NOMT**, naringenin 7-O-methyltransferase.

**Supplemental Figure S8.** Expression of diterpene phytoalexin biosynthetic genes in leaves of control and high-Fe plants (-, mock-inoculated; +, *M. oryzae*-inoculated). Transcript levels were determined by RT-qPCR analysis at 48 hpi. The expression values were normalized to the rice *Ubiquitin1* gene. Three independent biological replicates (2 technical replicates each) were assayed. Data are mean ± SEM (n=3). Asterisks indicate statistical significant differences calculated by two-way ANOVA (*, **, and *** indicate p < 0.05, 0.01, and 0.001, respectively). Gene-specific primers are listed in **Supplemental Table S8**.

**Supplemental Figure S9.** Expression of genes involved in Fe homeostasis in leaves control and high-Fe plants (-, mock-inoculated; +, *M. oryzae*-inoculated). Transcript levels were determined by RT-qPCR analysis at 48 hpi. The expression values were normalized to the rice *Ubiquitin1* gene. Three independent biological replicates (2 technical replicates each) were assayed. Data are mean ± SEM (n=3). Asterisks indicate statistical significant differences calculated by two-way ANOVA (*, **, and *** indicate p < 0.05, 0.01, and 0.001, respectively). Gene-specific primers are listed in **Supplemental Table S8**.

**Supplemental Figure S10.** Malate dehydrogenase assay in total protein extracts or apoplast washing fluid obtained from control and high-Fe plants (mock-inoculated and *M. oryzae*-inoculated). Five biological replicates for each condition (with 10 plants per condition were analyzed. Representative results are presented.

