## Supplementary figures and images for "Iron treatment induces defense responses and disease resistance against *Magnaporthe oryzae* in rice"

### Supplemental Figure 1

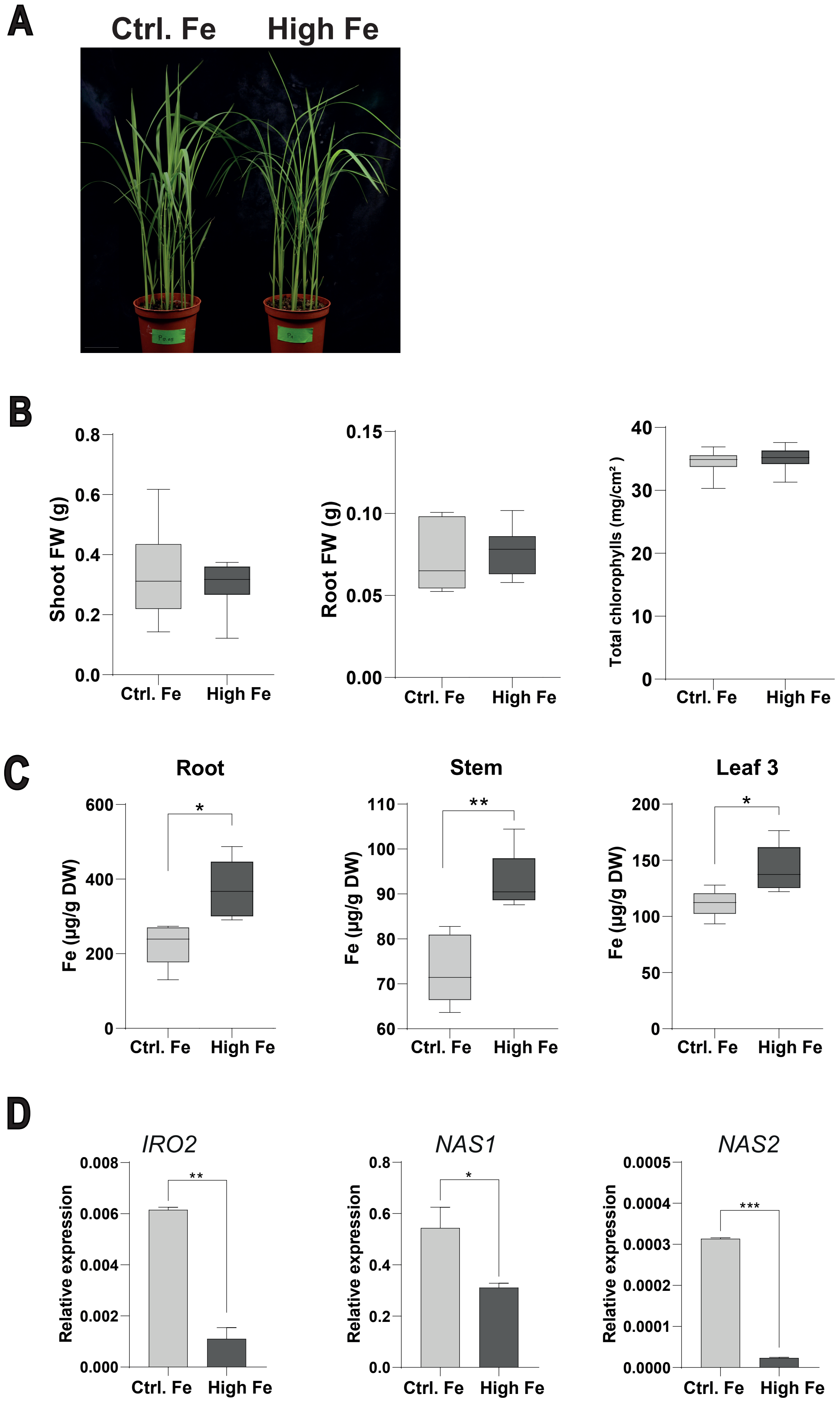

### Supplemental Figure 2

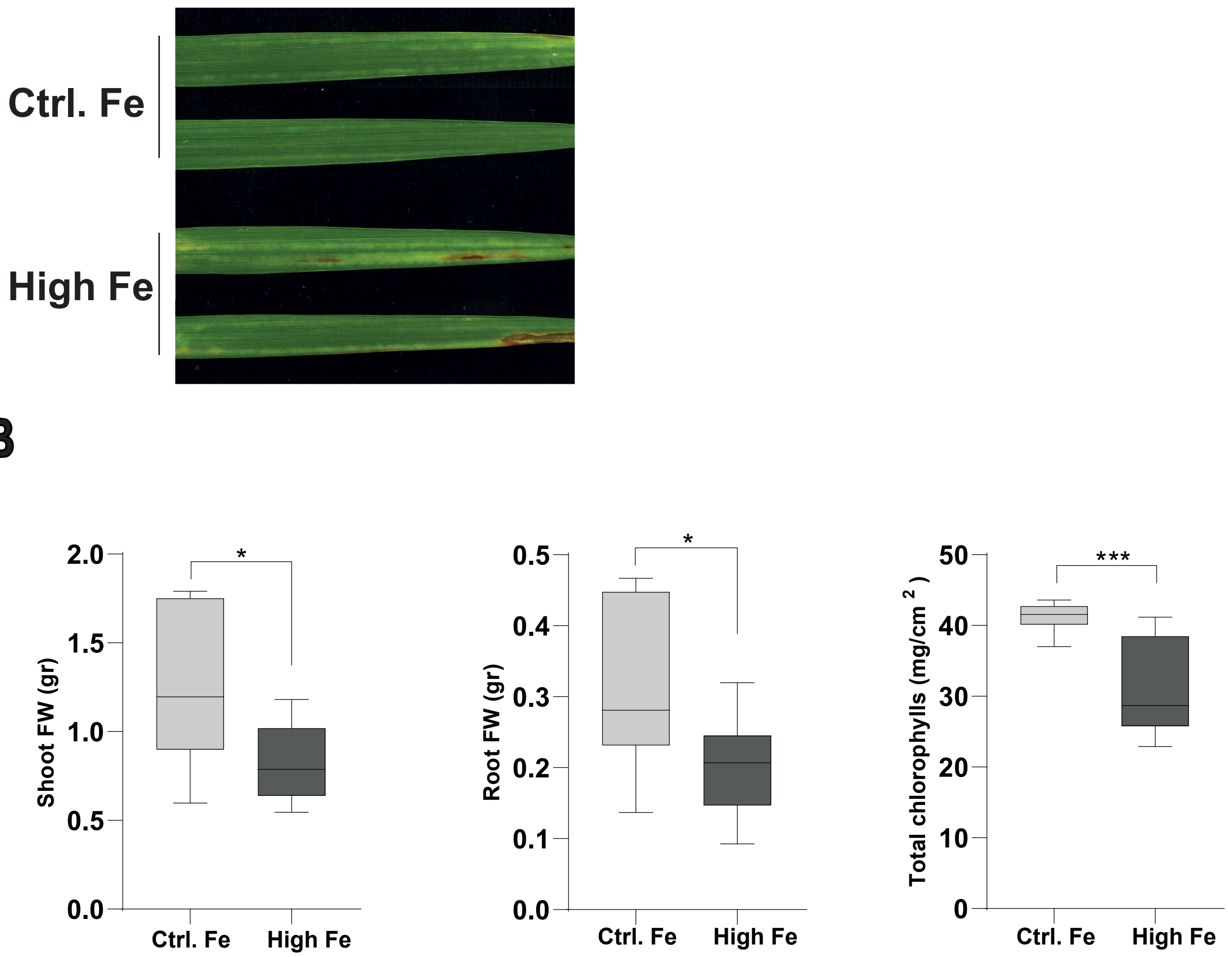

### Supplemental Figure 3

**High Fe**

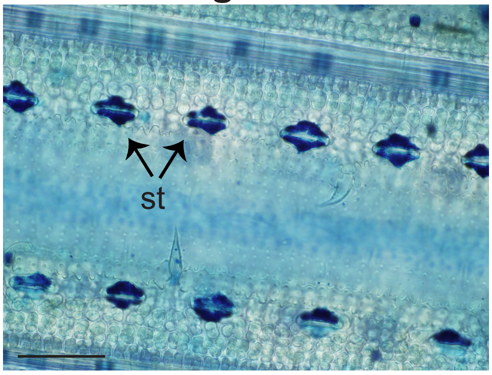

### Supplemental Figure 4

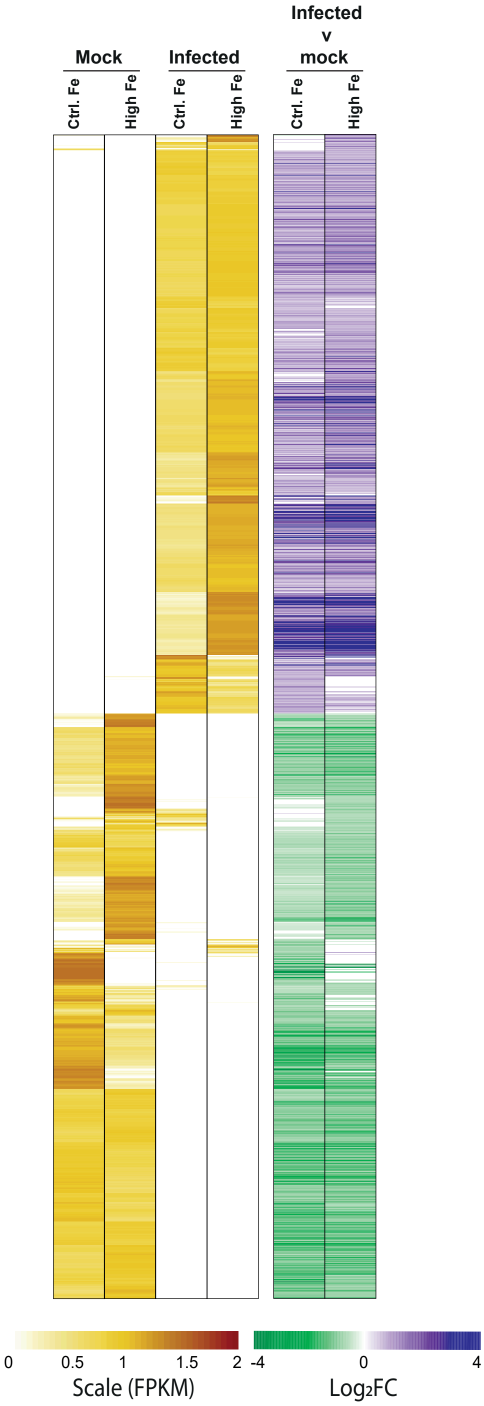

### Supplemental Figure 5

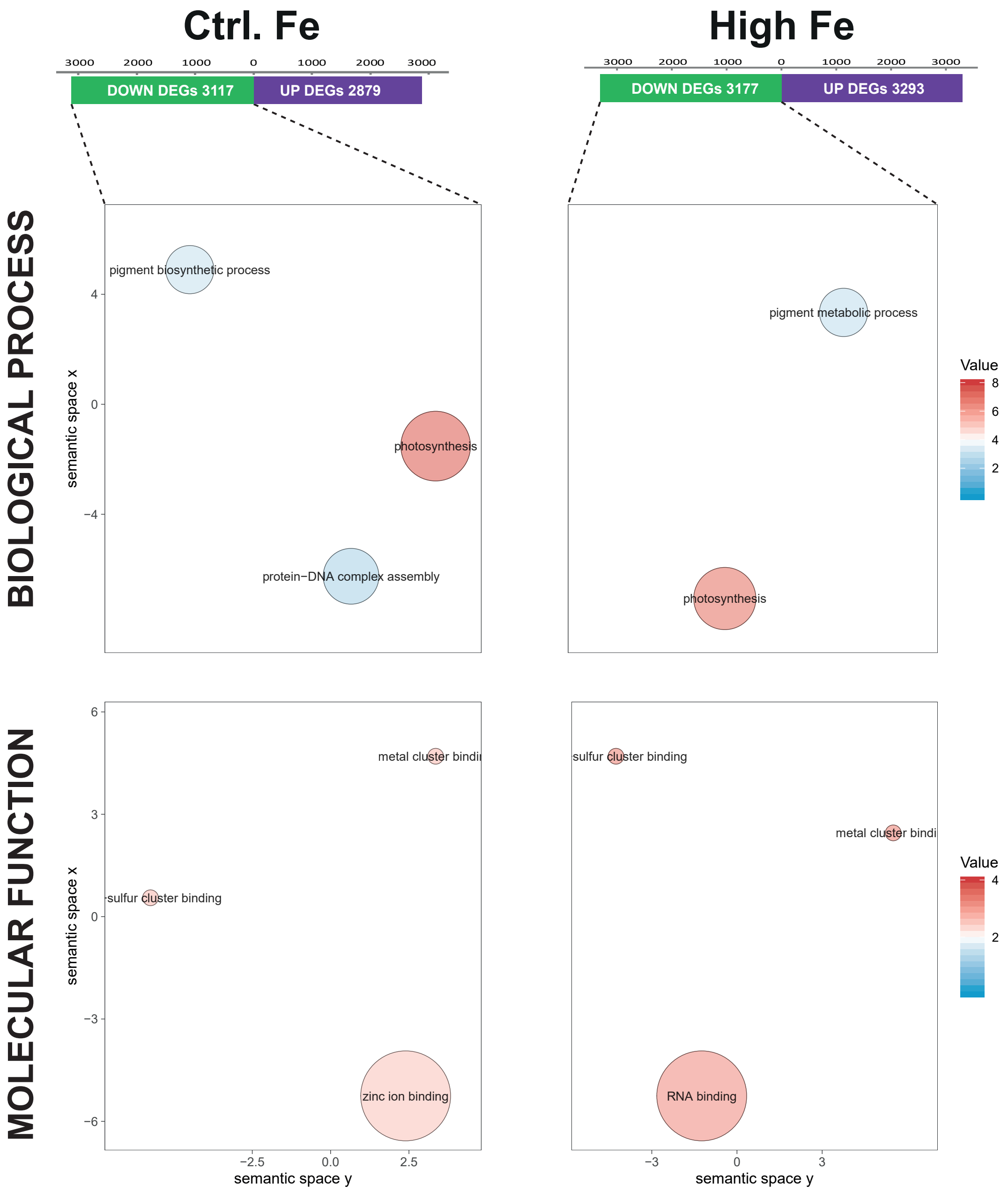

### Supplemental Figure 6

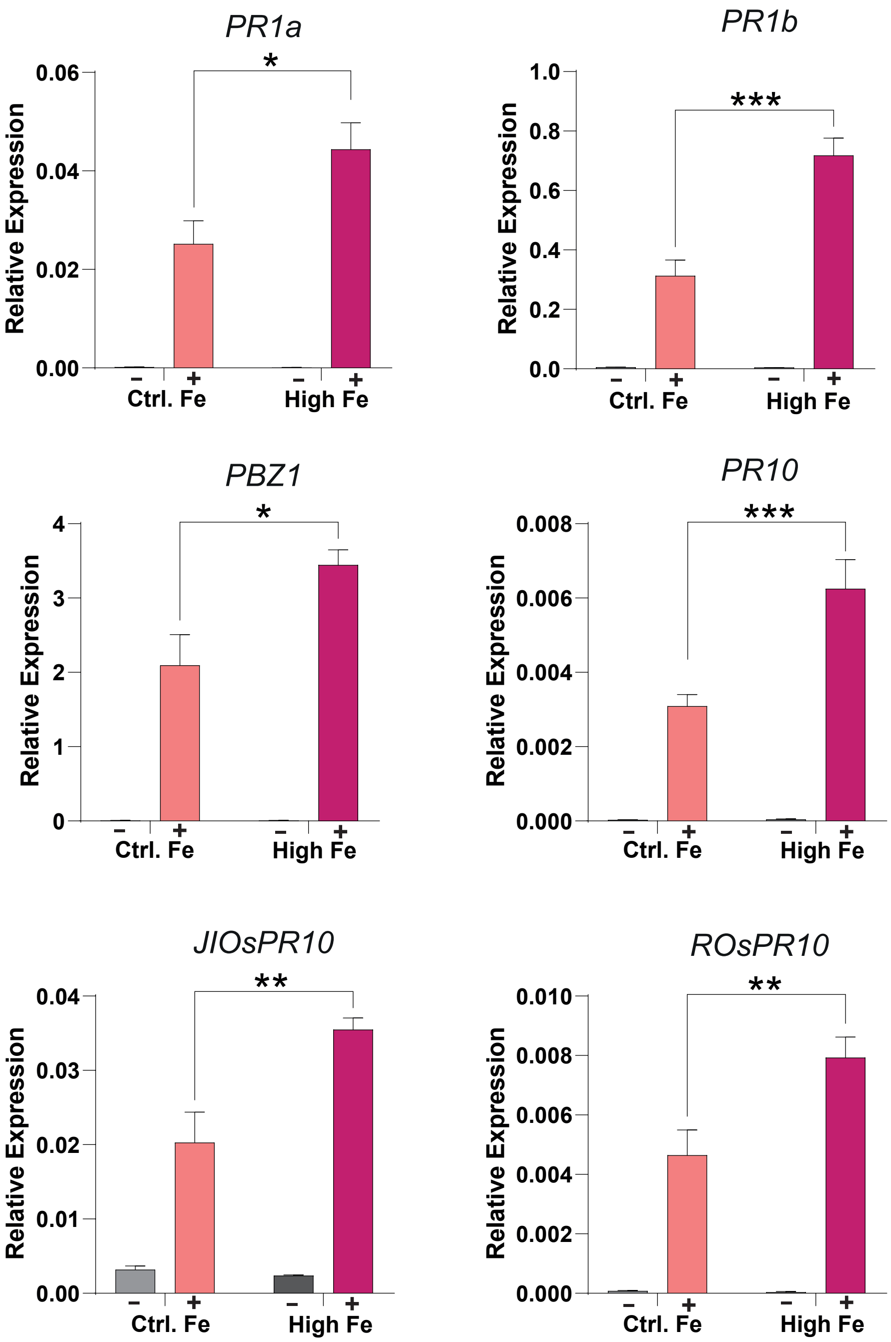

### Supplemental Figure 7

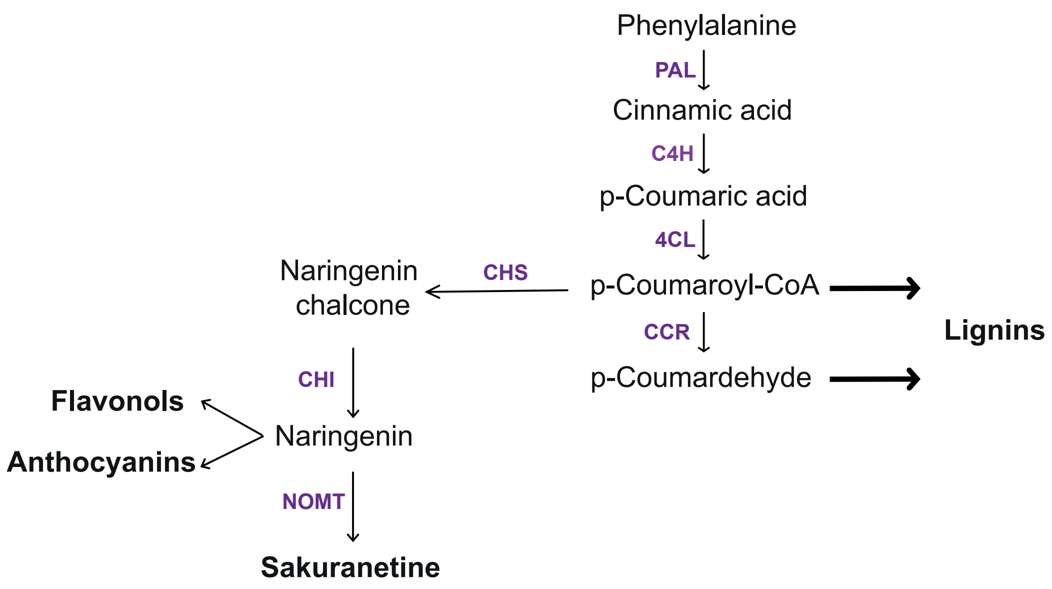

### Supplemental Figure 8

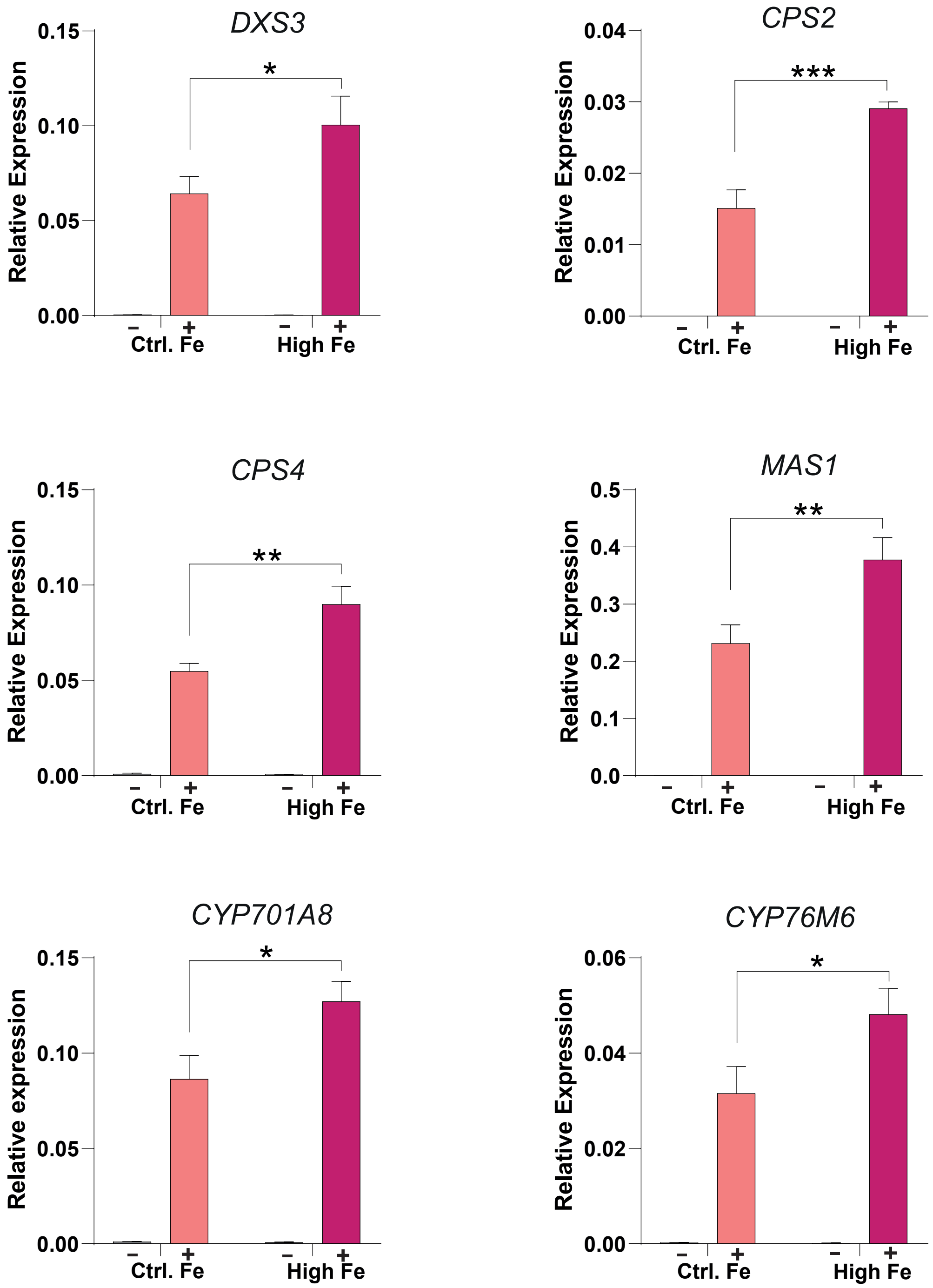

### Supplemental Figure 9

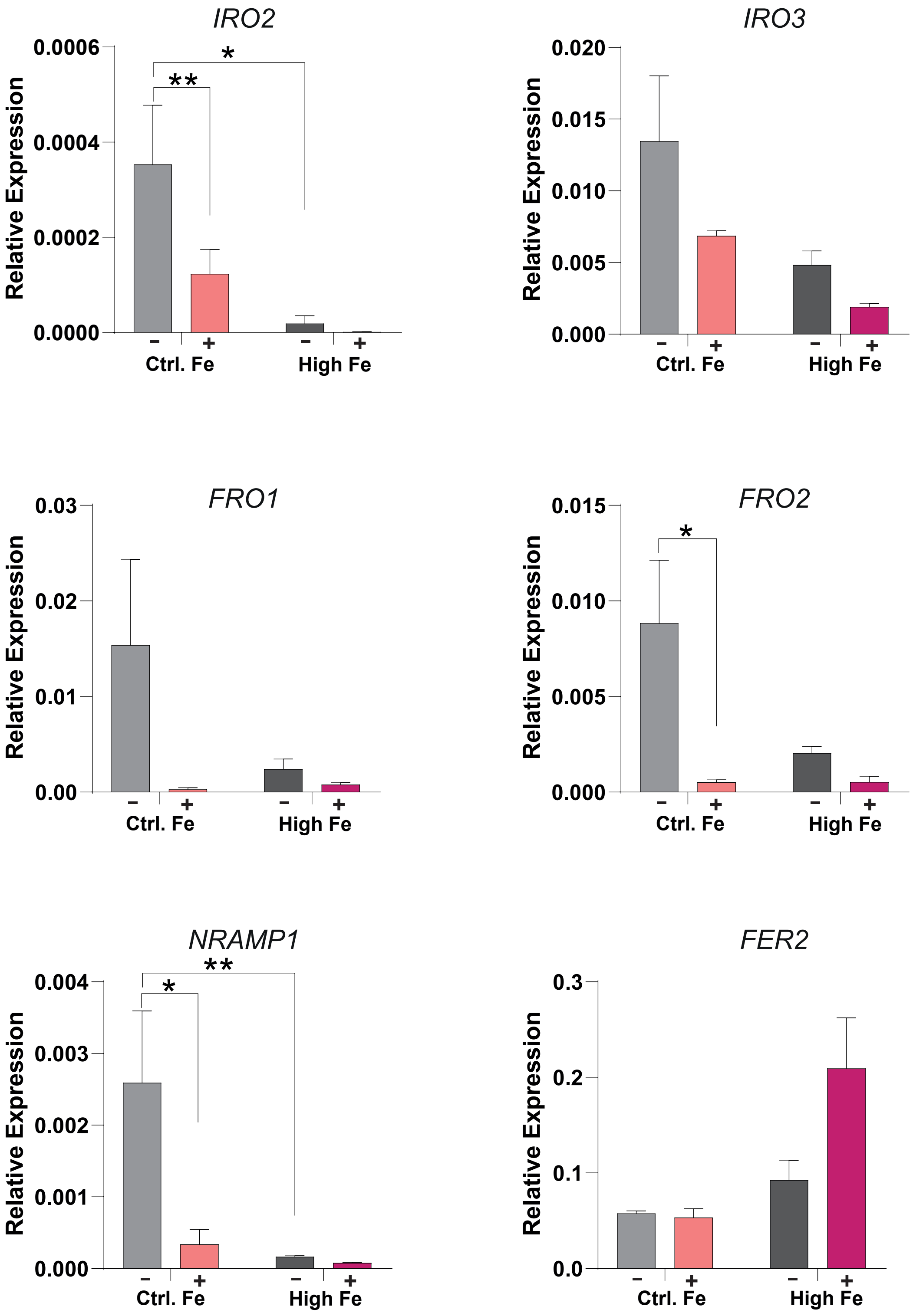

### Supplemental Figure 10

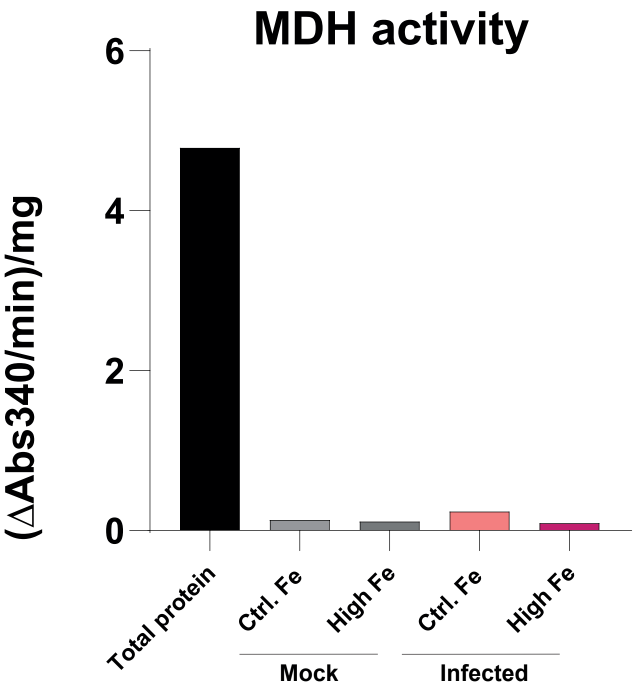
